# Trpv4 links environmental temperature to testicular differentiation in hermaphroditic ricefield eel

**DOI:** 10.1101/2025.06.09.658756

**Authors:** Yang Zhi, Luo Tingting, Zhang Yimin, Sun Yuhua

## Abstract

The ricefield eel (*Monopterus albus*), an economically important aquaculture species in China, is a freshwater teleost fish that exhibits protogynous hermaphroditism. Although progress has been made in understanding the sex determination and differentiation of this species, the underlying mechanisms remain unclear. Here we show that warm temperature promotes gonadal transformation by up–regulating testicular differentiation genes such as *dmrt1/sox9a* in ovaries. Trpv4, a Ca^2+^–permeable cation channel expressed in gonadal somatic cells, is highly sensitive to ambient temperature and links environmental temperature to testicular differentiation in ricefield eel. In female fish reared at cool temperature, injection of Trpv4 agonist into the ovaries leads to a significant up–regulation of male pathway genes, and in female fish exposed to warm temperature, Trpv4 inhibition or *trpv4* siRNA knockdown suppresses warm temperature–induced male gene expression. pStat3 signaling is downstream of Trpv4 and transduces Trpv4–controlled calcium signaling into the sex determination cascades. Inhibition of pStat3 activity prevents the up–regulation of testicular differentiation genes by warm temperature treatment and ovarian injection of Trpv4 agonist, whereas activation of pStat3 is sufficient to induce the expression of male genes, in the presence of Trpv4 antagonist. pStat3 binds and activates *jmjd3/kdm6b*, an activator of the *dmrt1* gene. Consistently, ovarian injection of Kdm6b inhibitor blocks the up–regulation of testicular differentiation genes by warm temperature exposure. We propose that environmental factors, such as temperature, promote gonadal transformation of ricefield eel by inducing the expression of male pathway genes in ovaries via the Trpv4–pStat3–Kdm6b–*dmrt1* axis. Our results provide new insights into the molecular mechanism underlying natural sex change of ricefield eel, which will be useful for sex control in aquaculture.

**Highlights:**

- Warm temperature promotes gonadal transformation of ricefield eel
- Trpv4 links environmental temperature and the sex determination pathway
- pStat3 is downstream of Trpv4-controlled calcium signaling
- pStat3 binds and activates *kdm6b/jmjd3*

## Introduction

Sex determination in animals is intriguing and fascinating. In mammals, sex is determined genetically (genotypic sex determination, GSD). In lower vertebrates such as fish and reptiles, however, sex regulators are diverse. Their sex can be influenced by various environmental factors, including temperature, pH, the breeding density and social status (Gutzke and Crews. 1988; Honeycutt et al. 2019; Mei and Gui, 2015; Todd et al., 2019), the so-called environmental sex determination (ESD). The temperature-dependent sex determination (TSD) is one of the best studied forms of ESD. In red-eared slider turtle (*Trachemys scripta*), American alligator (*Alligator mississippiensis*) and Atlantic silverside (*Menidia menidia*), sex is determined solely by the temperature during the thermosensitive period of embryogenesis. In Australian central bearded dragon (*Pogona Vitticeps*) and Nile tilapia (*Oreochromis niloticus*), which display GSD, temperature can override the genetic materials to control the gonadal sex differentiation (Deveson et al., 2017; Holleley et al., 2015). Analysis of expression data during embryogenesis of normal ZW females and temperature sex reversed ZZ females has provided important insights into temperature-driven sex determination in the bearded dragon (Whiteley et al., 2021). Irrespective of TSD or GSD+TE (temperature effects), the downstream components are fairly conserved, including the epigenetic factors such as *jmjd3*/*kdm6b* and sex determination genes such as *dmrt1* (Castelli et al., 2020; Lu et al., 2025; Martinez-Pacheco et al., 2024; Weber et al., 2020; Whiteley et al., 2020; Wu et al., 2024).

Most vertebrates, including the TSD reptiles, exhibit gonochorism. However, approximately 6% of fish species exhibit hermaphroditism, including protandrous, protogynous, and bidirectional hermaphroditism (Todd et al., 2016). The majority of them are marine fish, appearing in 27 families (Muncaster et al., 2013; Peng et al., 2020; Shao et al., 2014; Todd et al., 2019). Compared to marine fish, natural sex change in freshwater fish is very rare. The ricefield eel (*Monopterus albus*), also called Asian swamp eel, was firstly discovered as a protogynous hermaphroditic fish by Liu (Liu, 1944; Luo et al., 2026; Zhou and Gui, 2016). The species begins life as a female and then develops into a male through an intersex stage, thus displaying a female-to-male sex reversal during aging. Females are small in size (< 25 cm), and during and after sex change, there is a gradual increase in body size (> 55 cm for the majority of males). Among the described teleost fish species, ricefield eel has the fewest chromosome pairs (n= 12) with the fewest number of chromosome arms (Cheng and Zhou, 2022), and is emerging as an important model animal for studying sex determination and differentiation as well as adaptive evolution (Ji et al., 2001). As early as the Ming Dynasty in ancient China in 1578, pharmacist Shi-Zhen Li has described the medicinal value of ricefield eel in treatment of human diseases in his famous pharmacy monograph, the Bencao Gangmu, also called “Compendium of Materia Medica” (Cheng and Zhou, 2022). Nowadays, ricefield eel has been developed as one of the most important economical fish in freshwater aquaculture in China, with annual production exceeding 350,000 tons (Song et al., 2022). Unfortunately, the wild population has declined rapidly in the wild due to degradation of natural environment and human activity such as overfishing. The reproductive mode of ricefield eel, which leads to much more females than males in spawning season, severely affects the sex ratio, and decreases the productivity of broodstock. Moreover, adult females lay limited eggs (∼200) due to its small size, which is a limiting factor for massive production of seedling for aquaculture industry. Thus, the elucidation of the mechanisms underlying the sex determination/differentiation is important, which will aid in developing strategies/techniques for sex control that would break the bottleneck in aquaculture industry (Wu et al., 2019).

The life history of ricefield eel implies that environmental factors initiate and promote the gonadal transformation via epigenetic mechanisms. Consistently, histone demethyltransfearse/methyltransfearse genes such as *kdm6b/kmt2* and DNA methylation enzyme genes such as *dnmt1/3* were dynamically expressed throughout the sex change process, and the expression levels of the master sex determination/differentiation genes are closely correlated to the levels of DNA and histone methylation, which can be impacted by environmental exposure (Fan et al., 2021, 2022; Jiang et al., 2021; 2022; Hu et al., 2022; Wang et al., 2020). However, these epigenetic regulators are not inherently responsive to the environmental cues, implying that certain molecular sensors exist and serve as the link between environmental stimuli and the sex determination pathway. We have recently proposed that there is a temperature-induced sex reversal (TISR) mechanism in ricefield eel (Zhang et al., 2025), similar to that of embryonic bearded dragon (*P. Vitticeps*) (Whiteley et al., 2018). Importantly, isolated ovarian explants are responsive to temperature stimuli, suggesting that the perception of temperature is executed by certain sensors expressed in ovarian cells. While preliminary data have suggested that the Ca^2+^–permeable, non–selective cation channel Trpv4 (Transient Receptor Potential Vanilloid 4 channel) might be a potential thermosensor, how Trpv4-regulated signals are transduced into the sex determination cascades remains unclear (Zhang et al., 2025).

In this work, we hypothesized that Trpv4 may bridge environmental temperature and the sex differentiation pathway in ricefield eel. By using small molecule agonist and antagonist of Trpv4 as well as siRNA-mediated knockdown of *trpv4*, we provided solid evidences that temperature-evoked Trpv4 activity promotes the sex change of ricefield eel via the downstream Ca^2+^-pStat3-Kdm6b-*dmrt1* axis.

## Results

### Warm temperature promotes gonadal transformation

Natural populations of ricefield eel are mainly distributed across East and Southeast Asia. Previous studies have reported that at the onset of sex change, wild fish from different geographic populations and habitats vary in age, body weight and length. For instance, it is around 16 cm long (18-month-old) in Bandung area of Indonesia (Liem, 1963), and ∼20 cm in Hainan and Guangzhou areas of southern China (Chan and Philips, 1967; Wang and Zeng, 2006), ∼30 cm (2-year-old) in Wuhan area of central China (Wang et al., 2008), and 35-40 cm (3-year-old) in Tianjin area of northern China (Fan et al., 2017; 2021; Liu and Wang, 1987). The average annual temperature in Hainan, Guangzhou, Wuhan and Tianjin areas is approximately 25, 22, 17, and 13 □, respectively (Figure 1A). This observation implied that higher temperature facilitates the sex change of ricefield eel. To directly investigate this, during June-July, 2024, we have obtained ∼200 2-year-old wild ricefield eels from the southernmost Hainan and central Wuhan, and examined their gonads. We found that less than 5% of the fish from Wuhan area were intersex animals, whereas approximately 24% of the fish from Hainan area were in intersex stage (Figure 1B).

**Figure 1.**
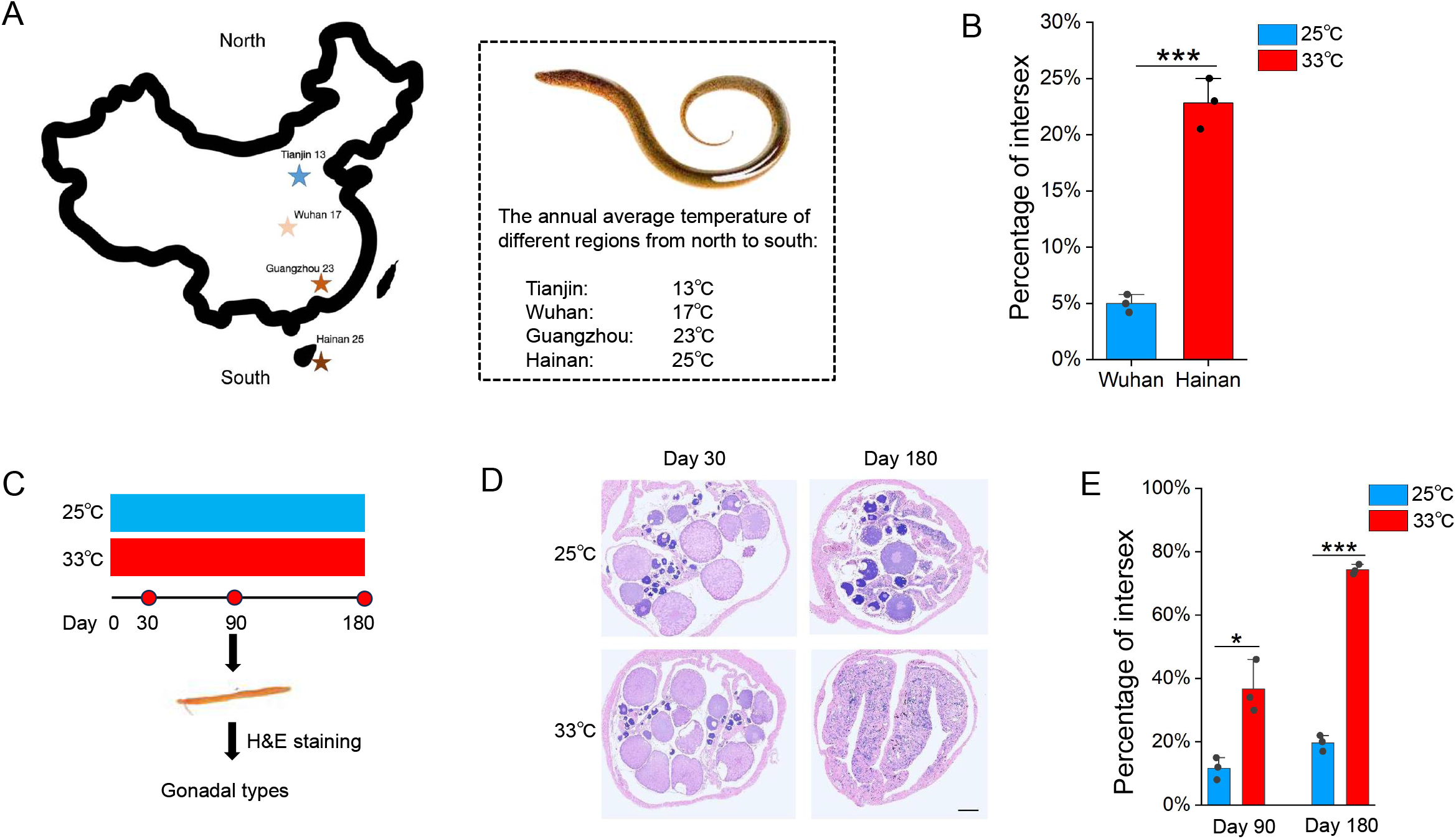
Warm temperature promotes gonadal transformation in ricefield eel. (A) The distribution of 4 geographic populations of ricefield eels in China, showing the average annual temperature of each. (B) Bar graph showing the percentage of intersex animals in 2-year-old wild-caught ricefield eels from Hainan (Hainan province) and Wuhan area (Hubei Province). n=200 for each group. (C) Diagram showing the design of long term temperature experiments. Two temperatures were used: 25 □ (cool temperature), 33 □ (warm temperature). n=400 for each group. (D) Representative H&E staining images showing the gonad types of animals that were reared at cool and warm temperatures at the indicated time points. Bar: 200 µm. (E) Bar graph showing the percentage of intersex animals after 180 days of cool and warm temperature treatment. The experiments were repeated at least two times.

In Wuhan area (Hubei province, China), the reproductive season of wild ricefield eel usually runs from May to July, when the average monthly temperature is 25-33 □ (Figure 1-figure supplement 1A). Immediately after spawning, the females (2-year-old) may undergo extensive ovarian tissue degeneration and physiological change (Liu and Gu, 1950), which leads to an irreversible commitment to becoming male via an intersex stage (Figure 1-figure supplement 1B-C). This observation again supported that the onset of sex change of ricefield eels is closely related to the external environment, in particular the warm temperature.

The above observations prompted us to hypothesize that warm temperature plays an important role in initiating and driving the sex change of ricefield eel. To directly test this, long term temperature experiments were performed using females from Wuhan Area (Figure 1C). One-and-a-half-year-old females (about 50 g) were randomly divided into two groups, and reared at 25/26 □ (cool temperatures, CT) and 33/34 □ (warm temperatures, WT), for a period of 6 months. At day 30, 90, and 180, the gonadal sex of randomly selected fish from different group was determined by H&E staining and expression analysis of sex-biased genes (Figure 1D). The average body length and weight of ricefield eels were comparable between the WT group and the CT group (Figure 1-figure supplement 1D-E). In CT group at day 90, ∼90% gonads were ovaries, and ∼10% were ovotestes. In WT group, however, ∼65% gonads were ovaries, and ∼35% were ovotestes (Figure 1E). In CT group at day 180, ∼80% gonads were ovaries, and ∼20% were ovotestes. In WT group, however, ∼25% gonads were ovaries, and ∼75% were ovotestes (Figure 1E). We concluded that warm temperature promotes gonadal transformation of ricefield eel.

### Trpv4 is highly responding to environmental temperature

We went on to investigate how the gonadal tissues are responding to temperature cues. Previous work has suggested that Trpv4 associates environmental temperature and sex determination in TSD alligator and ricefield eel (Huang et al., 2024; Yatsu et al., 2015; Zhang et al., 2025). We hypothesized that ricefield eel Trpv4 is expressed in ovary and functions as a thermosensor that perceives the environmental temperature cues. The results of qPCR experiments showed that *trpv4* was higher expressed in gonadal tissues than in non-gonadal tissues, exhibiting the highest expression in testis (Figure 2A). RNA in situ hybridization (ISH) experiments confirmed that *trpv4* levels increased from ovary to testis (Figure 2B), implying that it was functionally associated with testicular development during aging.

**Figure 2.**
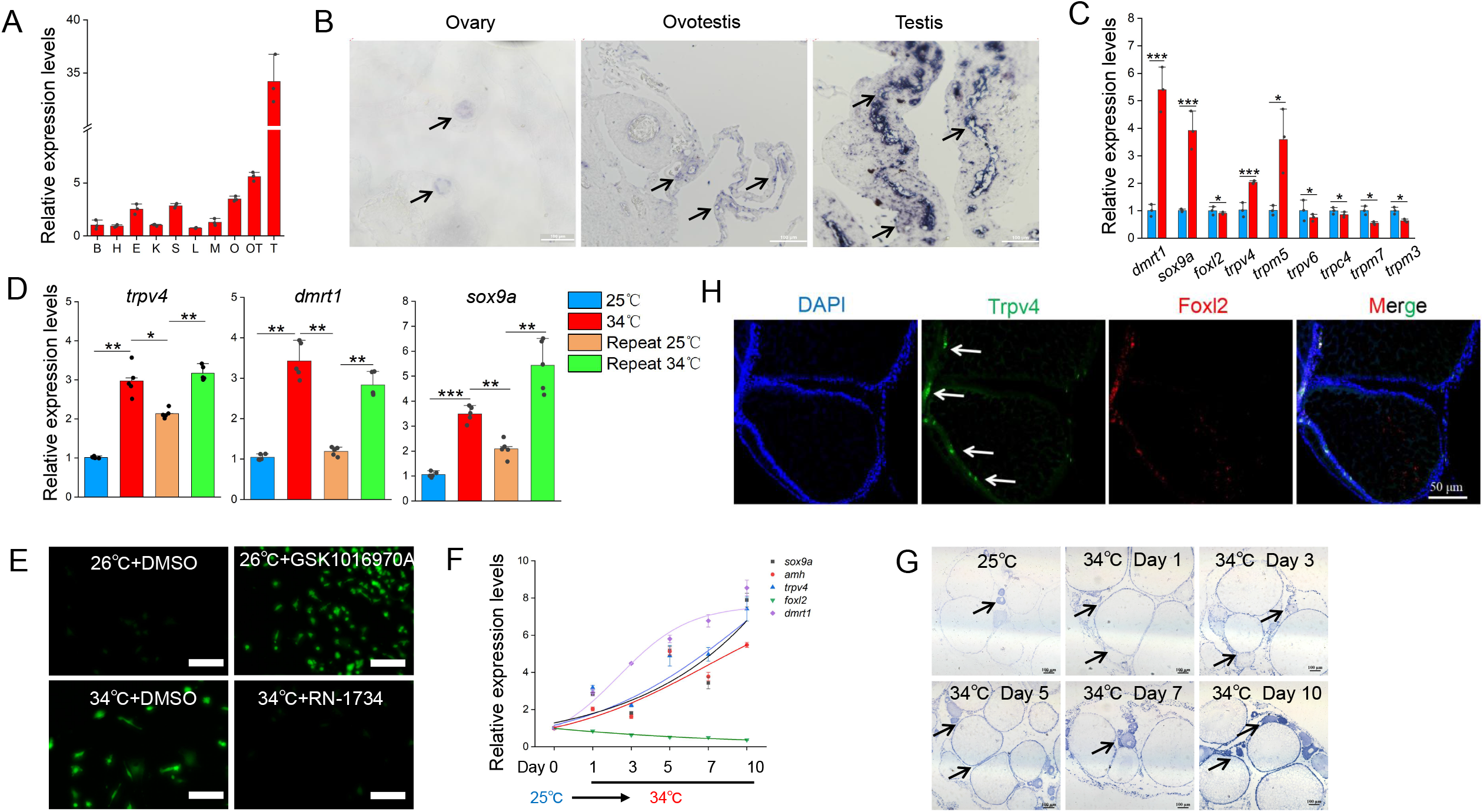
trpv4 is highly responding to environmental temperatures. (A) Relative expression levels of *trpv4* in 10 different tissues in adult ricefield eels. B: brain, H: heart, E: eye, K: kidney, S: spleen, L: liver, M:muscle, O: ovary, OT: ovotestis, T: testis. n=3. (B) ISH images showing *trpv4* expression in ovaries, ovotestes, and testes. Black arrows pointing to *trpv4* expressing cells. Bar: 200 µm. (C) The expression of the indicated *trp* and sex-biased genes of in vitro cultured ovaries at cool and warm temperatures. n=5 per group. (D) qPCR results showing the expression patterns of *trpv4* and male sex genes in repeated temperature shifting experiments of in vitro cultured ovaries. n=5 per group. (E) Confocal images showing the calcium signaling in cultured ovarian cells at the indicated conditions. After shifting to 34 □ for 0.5 hour, Cal-520 was added and calcium signal imaging was performed. The Trpv4 agonist GSK1016970A and antagonist RN1734 were administrated in the culture medium, and 15 minutes later, calcium signal imaging was performed. Bar: 100 µm. (F) qPCR results showing the dynamic expression of the indicated genes in gonads of female ricefield eel at 25 □, and at the indicated time points after shifting to 34 □. day 1: day 1 after shifting to 34 □. n=5 per group. (G) ISH images showing the dynamic expression of *trpv4* at the indicated time points before and after shifting to 34 □. At 25 □, *trpv4* was moderately expressed in follicles of various stages of developing oocytes, and interstitial cell types. After shifting to 34 □, *trpv4* signals became much stronger compared to 25 □. Bar: 200 µm. (H) Representative IF images showing the co-localization of Trpv4- and Foxl2-expressing cells. Bar: 50 µm. n=10. *: *P*< 0.05, **: *P*< 0.01, ***: *P*< 0.001, and ****: *P*< 0.0001. ns: not significant. All experiments were repeated at least three times.

In cultured primary ovarian explants, *trpv4* was one of the most up-regulated *trp* genes induced by warm temperature exposure (Figure 2C). Temperature-shifting experiments showed that *trpv4* was highly sensitive to temperature cues (Figure 2D), displaying a pattern similar to male sex genes such as *dmrt1/sox9a*, but inverse to that of female sex genes such as *cyp19a1a/foxl2* (Figure 2-figure supplement 1A). *trpv4* was elevated as early as 4 hours post warm temperature treatment (Figure 2-figure supplement 1B), suggesting that it was an early temperature-response gene. The *trpv4* gene encodes a constitutively active Ca^2+^-permeable cation channel that can be activated by warm temperature (Güler et al., 2002). Consistently, when Ca^2+^ indicator Cal-520 was added in the cells, a marked increase in the calcium signal was observed at 34 □ in comparison to 26 □ (Figure 2E), which suggested that Ca^2+^ signaling mediates the temperature-controlled Trpv4 activity, similar to the embryonic dragon (Whiteley et al., 2021). This was further supported by our RNA-seq data that *trpv4* and many genes involved in Ca^2+^ transport and sequestration were up-regulated in ovotestes compared to ovaries (Zhang et al., 2025).

While *trpv4* is highly responding to temperature changes in cultured ovarian cells, it is not known whether this was the case in vivo. To explore this, female fish were transiently reared at cool temperature (25 □) for 1 day, and then shifted to warm temperature (34 □) for 10 days. *trpv4* mRNA in ovaries was already elevated on day 1 after shifting to 34 □ (Figure 2-figure supplement 1C), and its levels progressively increased over time, exhibiting a pattern similar to that of testicular differentiation genes (Figure 2F). To understand in more detail of the role of *trpv4*, we studied its expression pattern in ovaries by performing ISH (mRNA in situ hybridization) experiment. The ISH results showed that *trpv4* was moderately expressed in ovarian somatic and interstitial cells around the oocytes at cool temperature (Figure 2G). Warm temperature exposure led to an increase of *trpv4* gene expression in ovarian follicles of different developmental stages, and its levels were positively correlated with the duration of temperature exposure.

To explore the identity of the Trpv4-expressing follicle cells, double immunofluorescence (IF) experiments were performed using antibodies against Trpv4 and Foxl2, a granulosa cell marker. The IF results showed that Trpv4 protein was expressed in a portion of Foxl2-positive granulosa cells (Figure 2H). The observation suggested that these *trpv4-*expressing follicle cells in ovary may play an important role in gonadal transformation in response to warm temperature.

### Temperature-induced male gene expression depends on Trpv4

The co-upregulation of *trpv4* and male pathway genes prompted us to ask whether the thermosensor Trpv4 in ovaries function to transduce the temperature cues into the sex determination cascades. To investigate this, animal experiments were performed by injecting into ovaries with small molecules RN1734 and GSK1016790A, known Trpv4 specific antagonist and agonist, respectively (Liu et al., 2021). The temperature shifting experiments were performed, and females were divided into 4 groups: 25 °C+ DMSO; 25 °C+ GSK1016790A; 34 °C+ DMSO; 34 °C+ RN1734 (Figure 3A). 3-10 days post injection, the gonads were isolated and used for the subsequent analyses.

**Figure 3.**
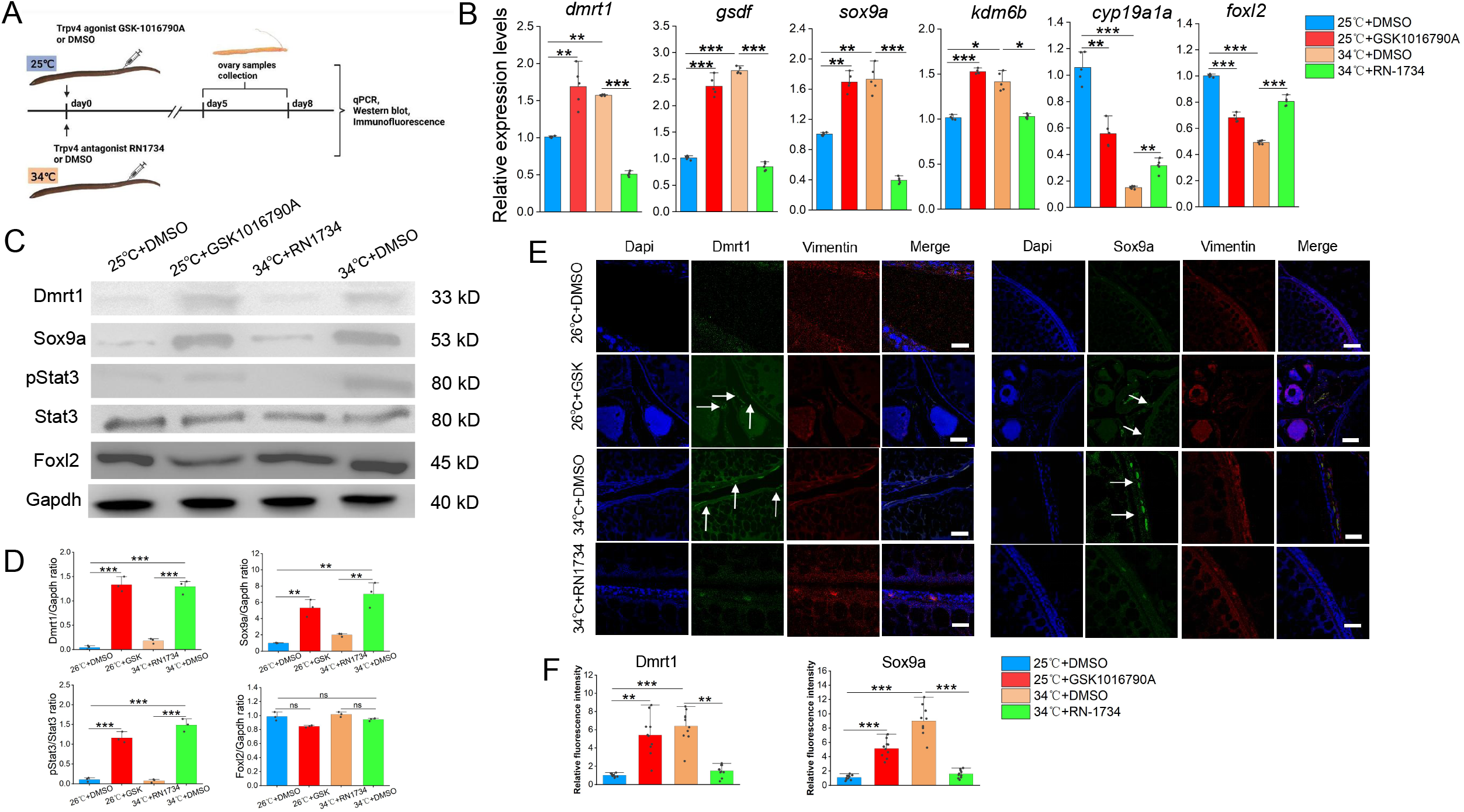
Warm temperature-induced male gene expression depends on Trpv4. (A) Cartoon showing the design of animal experiments. Female eels kept at cool (25 □) and warm (34 □) temperatures were injected with the Trpv4 agonist GSK1016790A and antagonist RN1734 into the ovaries, respectively. After 2-3 days of injection, the ovaries were isolated and processed for the subsequent experiments. n=40. (B) qPCR results showing the relative expression levels of the sex-biased genes at the indicated conditions, based on the animal experiments. n=5 per group. (C) Representative WB images showing the expression of the indicated markers at the indicated conditions. (D) Relative quantification of the indicated proteins in Panel C. WB were repeated 3 times. (E) Representative IF images showing the expression of the indicated markers at the indicated conditions. Vimentin was used for show all cell types in ovaries. GSK: GSK1016790A. Bar: 200 µm. n=10, and 9/10 showed induced expression of Dmrt1/Sox9a. (F) quantification of panel E. *: *P*< 0.05, **: *P*< 0.01, ***: *P*< 0.001, and ****: *P*< 0.0001. ns: not significant. All experiments were repeated at least three times.

Compared to the cool temperature group, warm temperature exposure increased the expression of testicular differentiation genes such as *dmrt1* and *gsdf*, accompanied by moderately decreased expression of ovarian differentiation genes such as *cyp19a1a* and *foxl2* (Figure 3B). Injection of 10 µM RN1734 into ovaries blocked the up-regulation of testicular differentiation genes by warm temperature treatment. Injection of 0.1 µM GSK1016790A in fish reared at 25 °C was sufficient to induce the expression of testicular differentiation genes to an extent similar to warm temperature treatment, accompanied by decreased expression of ovarian differentiation genes. The effects of small molecules on gene expression were dose dependent (Figure 3-figure supplement 1A). The effects RN1734 or GSK1016790A treatment on above sex determination factors were confirmed at the protein levels, as revealed by Western blot (WB) and IF experiments (Figures 3C–F). The experiments were also repeated using cultured ovarian explants and/or cells, and similar results were observed (Figure 3-figure supplement 1B-F). Activation and inhibition of Trpv4 ion channel function by the small molecules was correlated with increased and decreased calcium signaling, respectively (Figure 2E), suggesting calcium signaling mediated Trpv4 in the control of sex-biased gene expression.

The above data suggested that Trpv4 is sufficient and necessary to induce the expression of testicular differentiation genes in ovaries, via its ion channel function. To further demonstrate this, *trpv4* siRNA was injected into the ovaries. *trpv4* expression was markedly down-regulated by 0.1 µM siRNA injection (Figure 4A). Importantly, siRNA injection led to a marked decrease in expression of male pathway genes that were up-regulated by warm temperature exposure, and a slight increase in expression of ovarian differentiation genes that were repressed by warm temperature treatment. Similar results were observed at the protein levels as revealed by WB experiments (Figures 4B-C). We concluded that warm temperature-induced male gene expression is dependent on Trpv4 activity.

**Figure 4.**
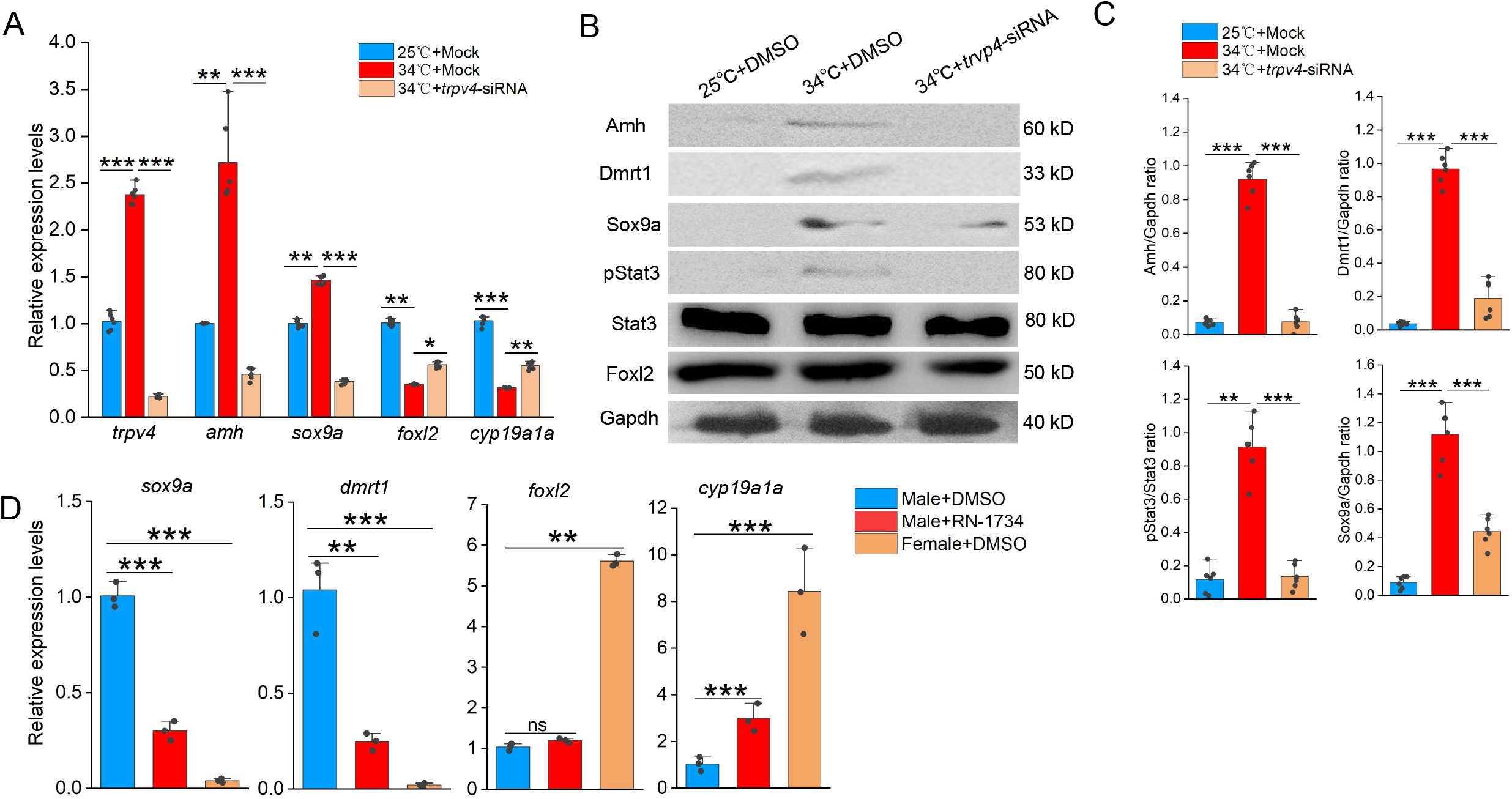
siRNA-mediated trpv4 knockdown abolishes the abnormal up-regulation of male genes by warm temperature treatment. (A) qPCR results showing the relative expression of the indicated genes at the indicated conditions in animal experiments. (B) Representative WB images showing the expression of sex biased proteins at the indicated conditions in animal experiments. (C) Quantification of panel B. (D) qPCR results showing the relative expression of the indicated genes at the indicated conditions. 10 µM RN1734 and DMSO was injected into gonads of male fish reared at 25/26 °C. Female fish injected with DMSO were used as control. *: *P*< 0.05, **: *P*< 0.01, ***: *P*< 0.001, and ****: *P*< 0.0001. ns: not significant. All experiments were repeated at least three times.

Next, we evaluated the effect of Trpv4 inhibition in testes of male adults. To this end, 10 µM RN1734 was injected into gonads of male fish, and qPCR was performed to assess the expression of female– and male pathway genes. The results showed that Trpv4 inhibition by RN1734 treatment led to an increase in the expression of female pathway genes in testes, at the expense of male pathway genes (Figure 4D). The results suggested that Trpv4 activity is persistently required for the expression of male pathway genes in testis.

### pStat3 signaling is downstream of Trpv4

We next asked how Trpv4-controlled Ca^2+^ flux was interpreted and transduced into the sex determination cascades. In TSD turtles and TISR central bearded dragon, Stat3/4 are activated through phosphorylation by temperature-evoked calcium signaling, and is directly involved in sex determination (Weber et al., 2020; Whiteley et al., 2021). Based on our RNA-seq data, we found that *stat3*, and the Jak/Stat3 pathway target genes such as *socs3* and *egr1/2*, were significantly up-regulated in early ovotestes compared to ovaries (Figure 5A). We therefore hypothesized that pStat3 signaling is downstream of Trpv4-contolled calcium influx to promote the expression of male pathway genes in ovaries. In fact, several lines of evidences were in favor of this hypothesis. First, activation of Trpv4 ion channel by GSK1016790A at cool temperature led to elevated pStat3 levels and calcium signals in ovarian explants, similar to that by higher temperature treatment, and inhibition of Trpv4 by RN1734 at warm temperature decreased pStat3 levels and calcium signals (Figures 5B-D). Second, pStat3 signals were detected in the gonadal somatic cells around the oocytes and interstitial cells, analogous to that of *trpv4* (Figure 5B; Figure 2G). Third, the levels of pStat3 were gradually increased from ovaries to testes in wild-caught ricefield eels, along with the male sex promoting factors such as Amh (Figures 5E-F; Figure 5-figure supplement 1A-B). Fourth, WB blot analysis of A23187- or BAPTA-AM-treated ovarian cells showed that exposure to the ionophore A23187 increased phosphorylation levels of Stat3 at 25 °C, whereas chelatin of calcium with BAPTA–AM led to diminished activation of Stat3 (Figures 5G-H). Collectively, these observations strongly suggested that Trpv4 cell autonomously controls phosphorylation of Stat3 via the regulation of calcium signal in gonadal somatic cells.

**Figure 5.**
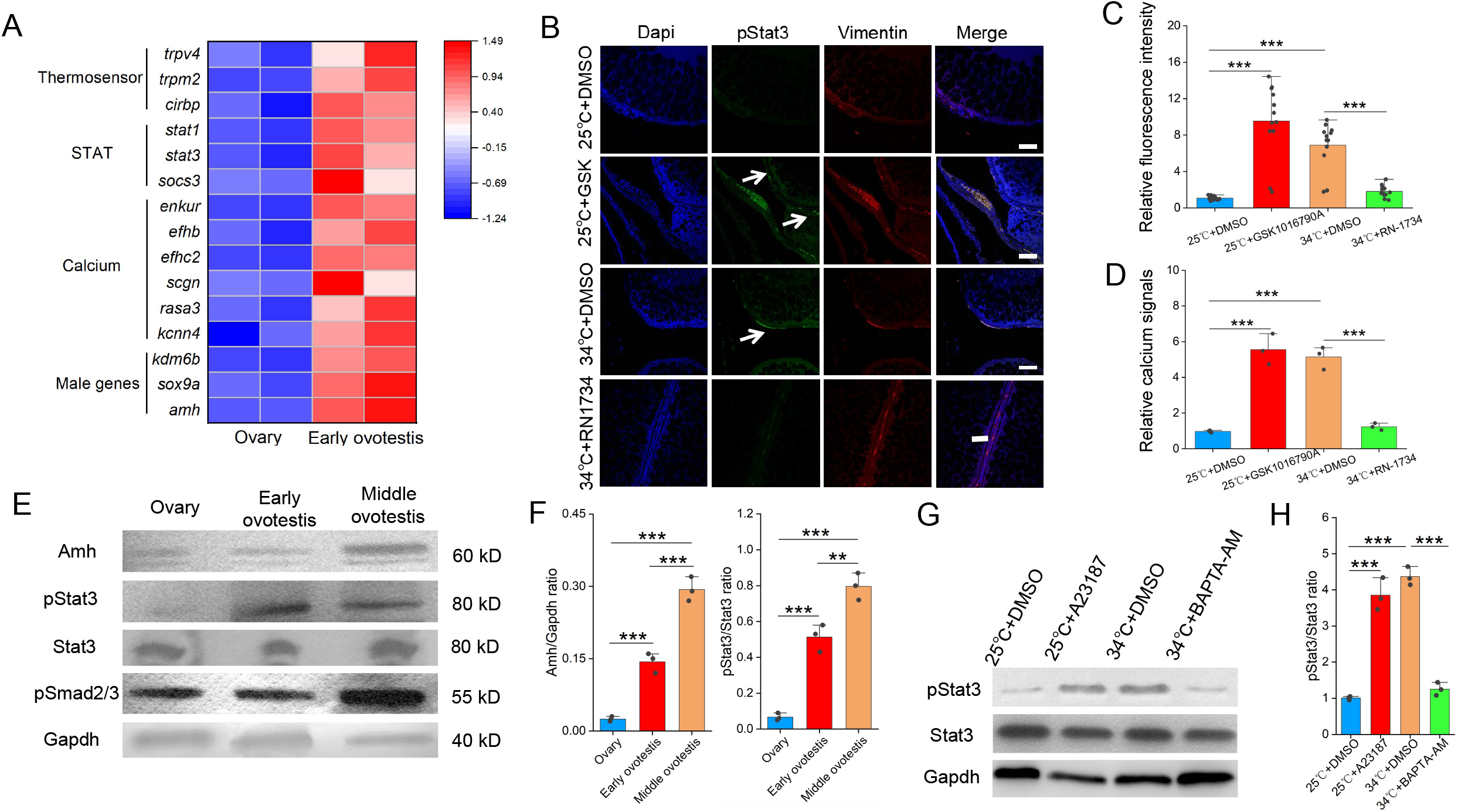
The JAK/Stat3 signaling is downstream of Trpv4. (A) Heat map showing the expression of the indicated genes of different groups. (B) IF images showing the expression of pStat3 at the indicated conditions in animal experiments. The white arrows indicated the location of pStat3 expressing cells. The experiments were repeated at least two times. Bar: 200 µm. n=12, and 10/12 showed increased expression of pStat3. (C) Quantification of panel B. (D) Bar graph showing the relative calcium signals at the indicated conditions. Ovarian explants were cultured at the indicated conditions, and calcium signals were determined by calcium indicator dye Cal-520 acetoxymethyl ester. (E) Representative WB images showing the expression of the indicated makers in ovaries, early ovotestes, and middle ovotestes. n=5 per group. (F) Quantification of panel E for relative expression of Amh and pStat3. (G) pStat3 levels after the addition of A23187 and BAPTA-AM in cultured ovarian cells at 25 °C and 34 °C conditions. (H) Quantification of panel G. All experiments were repeated at least two times.

To explore the function of the pStat3 signaling, animal experiments were performed by injecting into ovaries with small molecules HO-3867 or Colivelin. HO-3867, a curcumin analogue, is a selective pStat3 inhibitor, which blocks pStat3 activity by directly binding to Stat3 DNA binding domain, and Colivelin is a potent synthetic peptide activator of pStat3, which increases pStat3 levels by acting through the GP130/IL6ST complex (Wu et al., 2024). The injected females were divided into 4 groups based on the temperatures and the small molecules injected: 25 °C+ DMSO; 25 °C+ Colivelin; 34 °C+ DMSO; 34 °C+ HO-3867 (Figure 6A). 3-10 days post injection (dpj), the ovaries were isolated and subjected to the subsequent analyses. Injection of HO-3867 blocked the up-regulation of male pathway genes by warm temperature exposure, whereas injection of Colivelin at 25 °C was sufficient to induce the expression of these genes to an extent similar to warm temperature treatment (Figures 6B). Similar results were observed at the protein levels as revealed by IF experiments (Figures 6C-D). We also repeated the experiments using ovarian explants and/or cells, which produced similar results (Figure 6–figure supplement 1A-B).

**Figure 6.**
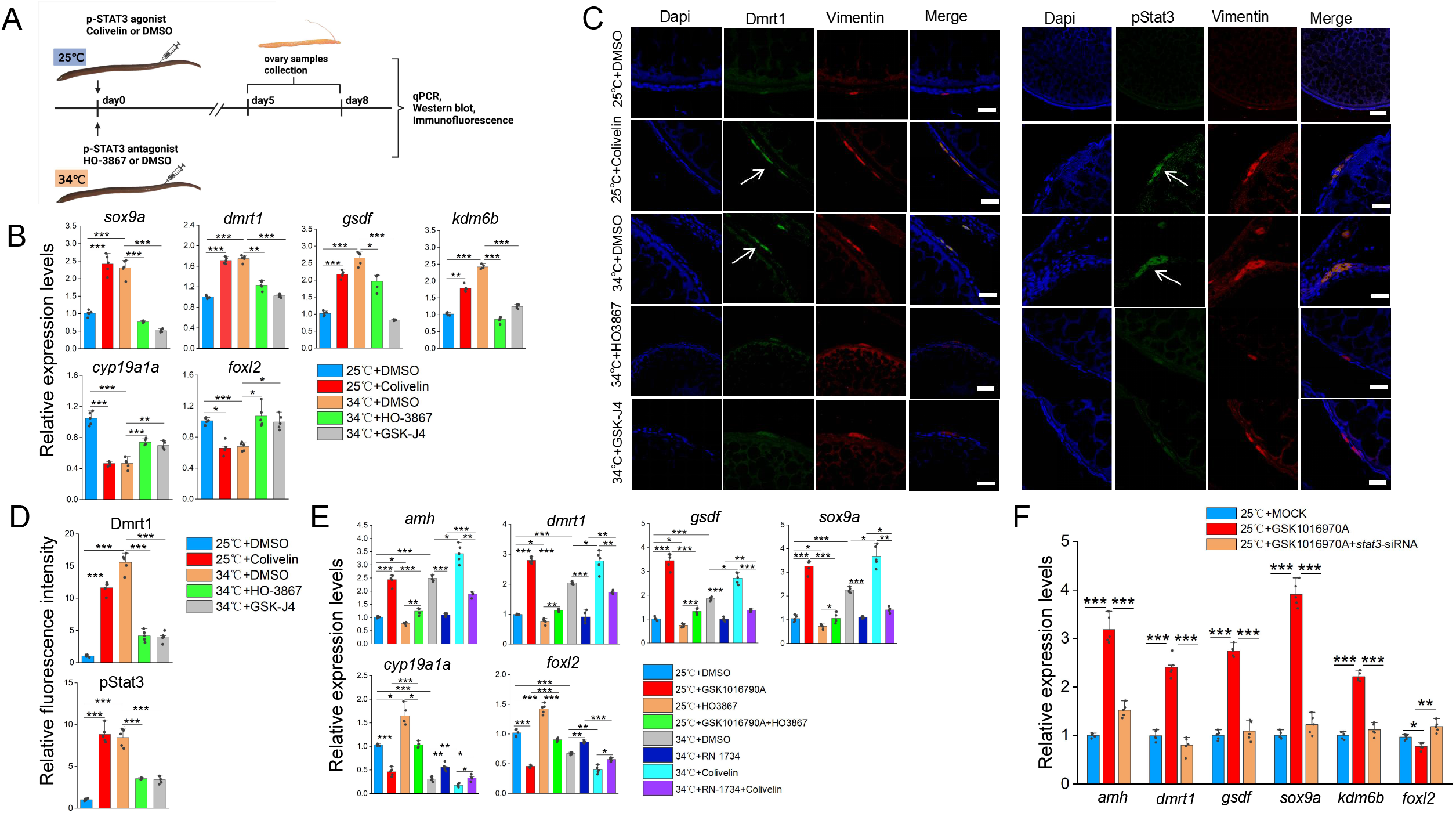
Animal experiments showing that warm temperature induced male gene expression depends on pStat3. (A) Cartoon showing the design of the experiments. Female eels kept at cool (25 □) and warm (34 □) temperatures were injected with the pStat3 agonist Colivelin and antagonist HO3867 into the ovaries, respectively. After 2-3 days of injection, the ovaries were isolated and processed for the subsequent experiments. n=50. (B) qPCR results showing the relative expression of the indicated genes at the indicated conditions. n=5 per group. (C) IF images showing the expression of male biased genes at the indicated conditions. Bar: 200 µm. n=10, and 8/10 showed increased expression of pStat3/Dmrt1. (D) Quantification of panel C. n=5 per group. (E) qPCR results showing the relative expression of the indicated genes at the indicated conditions. n=5 per group. (F) qPCR results showing the relative expression of the indicated genes at the indicated conditions. n=5 per group. *: *P*< 0.05, **: *P*< 0.01, ***: *P*< 0.001, and ****: *P*< 0.0001. ns: not significant. The qPCR experiments were repeated at least three times.

To functionally demonstrate that pStat3 signaling is downstream of Trpv4, rescue experiments were performed by injecting into ovaries with individual and combined small molecules as described below. The females were divided into 6 groups based on the temperatures and the small molecules injected: 25 °C+ DMSO; 25 °C+ GSK1016790A; 25 °C+ GSK1016790A+HO-3867; 34 °C+ DMSO; 34 °C+ RN1734; 34 °C+ RN1734+ Colivelin. 3-5 days post injection, the ovaries were isolated and subjected to the qPCR analysis. The results showed that mRNA up-regulation of testicular differentiation genes by the administration of Trpv4 agonist at 25 °C was abolished by the treatment with pStat3 inhibitor, and that expression of testicular differentiation genes inhibited by Trpv4 antagonist treatment at 34 °C can be partially restored by the injection with pStat3 agonist (Figure 6E). Similar results were obtained using in vitro cultured ovarian explants (Figure 6-figure supplement 1C).

To further confirm that Trpv4 functions upstream of pStat3, Trpv4 agonist GSK1016790A and *stat3* siRNA were simultaneously injected into the ovaries of fish reared at 25 °C, and the expression of sex-biased genes was examined by qPCR analysis. The results showed that the testicular differentiating genes failed to be activated by GSK1016790A in the presence of *stat3* siRNAs (Figure 6F). Taken together, we concluded that Trpv4 promotes male sex gene expression in ovaries via the activation of the downstream pStat3 signaling.

### pStat3 binds and activates kdm6b

Kdm6b is known a conserved activator at the top of the male sex determination pathway, via binding and activating *dmrt1* through removing the repressive histone mark H3K27me3 (Chen et al., 2024; Dupont et al., 2025; Ge et al, 2018; Lu et al., 2025; Yao et al;, 2024), and pStat3 can transcriptionally activate the *kdm6b* gene by directly binding to its upstream DNA motifs in reptiles (Wu et al., 2024; Weber et al., 2020). We reasoned that pStat3 had a similar function in ricefield eel. When analyzing -5 kb promoter sequences upstream the TSS of the *kdm6b* gene, we found that there were three conserved pStat3 binding sites (TTCnnnGAA) (Figure 7A). Chromatin immunoprecipitation (ChIP) experiments using pStat3 antibodies showed that pStat3 levels were significantly higher at *kdm6b* promoter in ovaries of females reared at warm temperature than at cool temperature, which was abolished in the presence of pStat3 inhibitor HO-3867 (Figure 7B). To explore whether pStat3 directly activates *kdm6b*, luciferase assay was performed. pGL4-*kdm6b*-luc construct was constructed by cloning ∼3.5 kb *kdm6b* promoter sequences into the pGL4-luc plasmid. A mutant construct (pGL4-*kdm6b*M-luc) was generated by replacement of “TTCAGAGAA” with “TTAAAAGAA”. pGL4-*kdm6b*-luc and pGL4-*kdm6b*M-luc were transfected with into HEK293T cells in the presence and absence of pStat3 agonist Colivelin, and the luciferase activities were measured. The results showed that activities of pGL4-*kdm6b*-luc were significantly higher than that of pGL4-*kdm6b*M-luc in the presence of Colivelin (Figure 7C).

**Figure 7.**
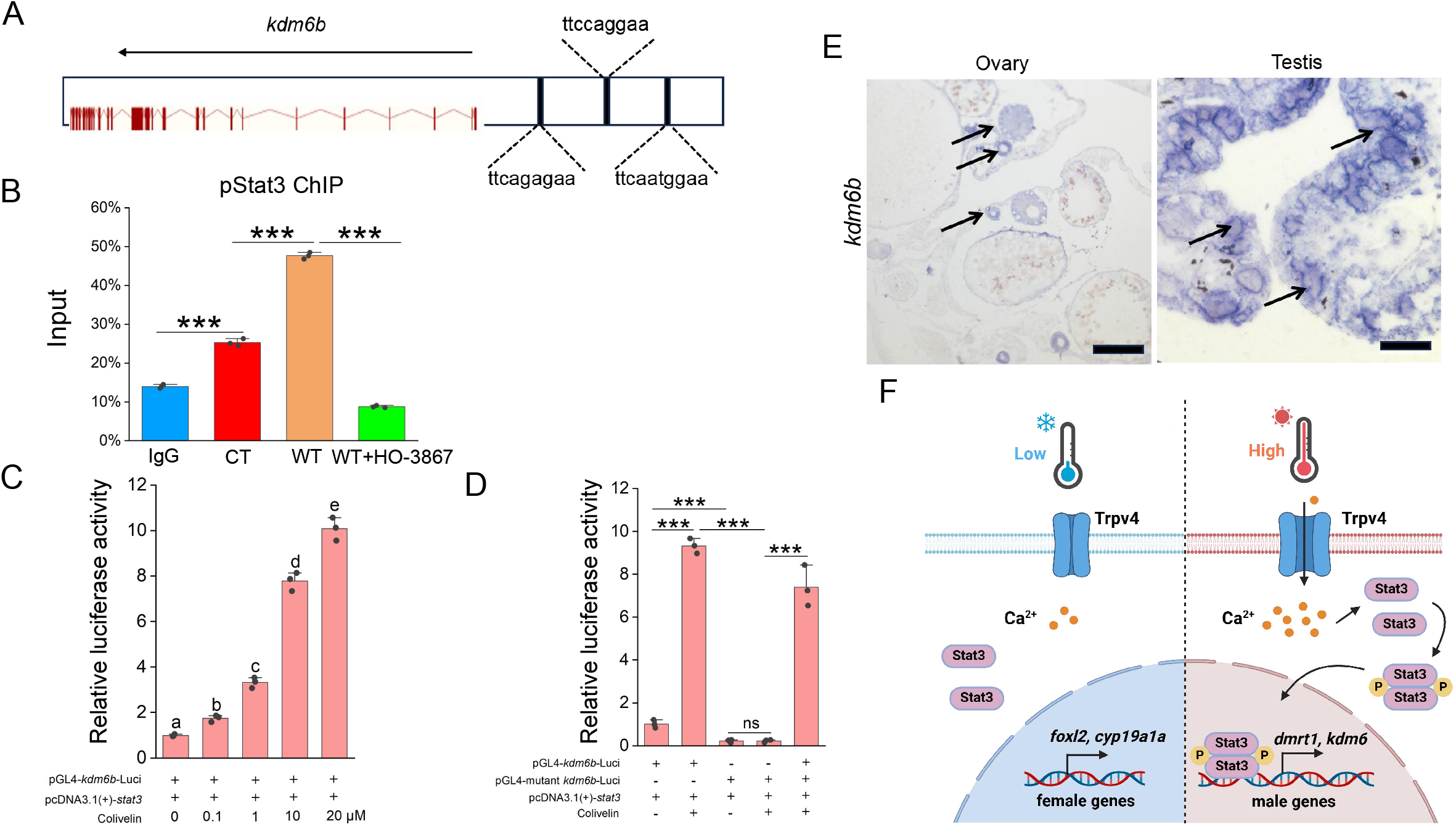
pStat3 binds and activates the kdm6b gene. (A) Cartoon showing the conserved pStat3 binding motifs upstream the TSS of the *kdm6b* gene. (B) ChIP experiments showing the enrichment of pStat3 at the *kdm6b* locus in ovarian tissues of fish reared at cool temperature (CT) and warm temperature (WT) conditions, in the absence and presence of HO-3867. (C) Luciferase assay for *kdm6b*-luc and *kdm6b*M-luc activities in 293T cells, in the absence and presence of pStat3 agonist Colivelin. (D) The representative ISH images showing the expression of *kdm6b* in ovary and testis. Bar: 200 µm. n=3 per group. (E) Cartoon showing how Trpv4 may link environmental temperature to the sex determination cascades via the downstream signaling pathways in ricefield eel. *: *P*< 0.05, **: *P*< 0.01, ***: *P*< 0.001, and ****: *P*< 0.0001. ns: not significant. ChIP and Luciferase experiments were repeated two times.

*kdm6b* mRNA was expressed in immature oocytes in ovaries, was induced by warm temperature exposure, and higher expressed in testes than in ovaries in ricefield eel (Figure 3-figure supplement 1B; Figure 6-figure supplement 1A; Figure 6B; Figure 7D), analogous to that of *trpv4*. The expression of *kdm6b* was elevated by the injection (animal experiments) or addition (cell culture experiments) of GSK1016790A and Colivelin at 25 °C, and was decreased by the injection or addition of RN1734 or HO-3867 at 34 °C (Figure 3B; Figure 6B; Figure 3–figure supplement 1B; Figure 6–figure supplement 1A). Thus, the *kdm6b* gene displayed an expression pattern similar to that of the testicular differentiation genes, which strongly supported that *kdm6b* is a male pathway gene downstream of Trpv4-Ca^2+^-pStat3 axis.

If *kdm6b* is downstream of Trpv4-pStat3 to regulate the expression of *dmrt1*, inhibition Kdm6b demethyltransferase activity should prevent up-regulation of testicular differentiation genes by warm temperature treatment, similar to HO-3867 and RN1734 treatment. 0.5 µM GSK-J4, an Kdm6b specific inhibitor, was injected into the ovaries of ricefield eel, and the expression of sex-biased factors were examined by the qPCR and IF analyses. The results showed that GSK-J4 injection significantly down-regulated the expression of testicular differentiation factors at the expense of ovarian differentiation factors (Figures 6B-D). Similar results were obtained in cultured ovarian cells (Figure 6-figure supplement 1A). We propose that there exists a Trpv4-pStat3-Kdm6b axis that links the environmental temperature and the male sex determination pathway (Figure 7E).

## Discussion

Since the first discover that ricefield eel is a teleost fish of hermaphroditism (Liu et al. 1944), the underlying mechanism has been under intensive investigation. However, it is still mysterious, partially because it is challenging to perform genetic studies due to unique reproductive strategy and life history of the species (Song et al., 2022). In this work, we provided solid evidences that warm temperature promotes the sex change of adult ricefield eel, and that the ion channel protein Trpv4 functions as an important molecular linker connecting the environmental temperature and the sex determination pathway. Our results support a model in which temperature-driven sex change is achieved via the Trpv4-pStat3-Kdm6b-Dmrt1 axis. In this model, warm temperature exposure leads to increased calcium influx via Trpv4 in ovarian granulosa cells, which promotes phosphorylation of pStat3 and expression of the male pathway genes, eventually and gradually resulting in transformation of ovary to testis via an ovotestis. Our work for the first time provides the mechanistic explanation of how natural sex change occurs in aging ricefield eel, which may serve as a paradigm to study natural sex change of other adult animals, including the hermaphroditic marine fish.

The initial perception and translation of environmental cues into the sex determination cascades remain unclear in any species. Previous studies have proposed endoplasmic reticulum chaperone, heat shock proteins and transmembrane ion channels, as potential temperature sensors (He et al., 2010; Shi et al., 2024). In contrast to Cdc2-like kinases (Clks) that response to subtle temperature change in homeothermic organisms, the transient receptor potential (TRP) cation channels may play important roles in poikilothermic animals that are exposed to strong ambient temperature change (Haltenhof et al., 2020; Huang et al., 2024). The TRP channel family can be divided into at least seven subfamilies, including TRPA, TRPC, TRPM, TRPML, TRPN, TRPP and TRPV. The thermo-TRPs contain nine members, such as TRPV (TRPV1-4). TRPV1-3 are mainly activated by temperatures above 35 °C, respectively (Mo et al., 2022). Of note, TRPV4 is activated by warm temperatures (27-35 °C) (Goikoetxea et al., 2021; Güler et al., 2002; Fujita et al., 2017), which are physiologically relevant to the spawning and/or the onset of sex change of ricefield eel. Trpv4 has been shown to be abundantly expressed during testicular/sperm development (Kumar et al., 2016; Mundt et al., 2018). Activation of Trpv4 by various endogenous/exodogenous stimuli increases Ca^2+^ influx, participating in multiple downstream biological events, including gonadal homeostasis (Guler et al., 2002; Liu et al., 2021; Luo et al., 2023; Vrenken et al., 2016; Yamamoto et al., 2024). Importantly, Trpv4 has been shown to associate temperature and sex determination in TSD alligator, and pharmaceutical activation and inhibition of Trpv4 alters testis differentiation (Yatsu et al., 2015). In ricefield eel, *trpv4* was one of the most up-regulated *trp* genes by warm temperature treatment, and its expression was associated with testicular development. Thus, we focused on the role of Trpv4 in this work. Based on in vitro and in vivo experiments, we demonstrated that a portion of Trpv4 expressing gonadal somatic cells are highly sensitive to temperature cues and are functionally important in linking environmental temperature to sex determination pathways in ricefield eel.

Previous work in the red-eared slider turtle (*Trachemys scripta elegans*) and the bluehead wrasse (*Thalassoma bifasciatum*) has indicated that Stat3 phosphorylation and epigenetic regulation are involved in TSD or sex change (Todd et al., 2019). In turtles, pStat3 is likely to be involved in sex determination by transcriptionally activating the female pathway genes such as *foxl2* and/or repressing the epigenetic factors as *kdm6b* (Chen et al., 2024; Deveson et al., 2017; Ge et al., 2018; Holleley et al., 2015; Weber et al., 2020; Wu et al., 2024). The function of *jmjd3/kdm6b* in sex determination in embryonic TISR fish such as tilapia has been well established (Yao et al., 2023). Overexpression and knockdown of *kdm6b* cause female to male or male to female sex reversal, respectively. Our ChIP and luciferase reporter experiments suggested that pStat3 directly binds and activates the *kdm6b* gene in ricefield eel. In turtles, pStat3 functions as a repressor of *kdm6b* (Weber et al., 2020), whereas in ricefield eel and the bluehead wrasse, pStat3 exerts an opposite effect (Todd et al., 2019). We reasoned that a yet-unidentified co-factor may determine whether Stat3 is a transcriptional repressor or activator. A comparison of promoter sequences of *kdm6b* between turtle and ricefield eel supported this (data not shown).

Trpv4, pStat3 and Kdm6b were expressed in somatic cells surrounding ovarian oocytes and interstitial cells. After warm temperature exposure, their levels were elevated over time, similar to that of the male pathway genes. We propose that the magnitude and duration of temperature exposure promote sex change of ricefield eel by driving the accumulation of testicular differentiation genes in sufficient quantities (Weber et al., 2020). In sex reversal animals, a long existing question is what cells in the gonad respond to environmental temperatures to initiate sex change. In groupers such as *Epinephelus akaara*, certain pre-existing somatic cells, called testicular-inducing steroidogenic cells, trigger the sex change (Murata et al., 2021). Recently, it has been proposed that a group of Nr5a1^+^Trp^-^ thermosensitive steroidogenic cells use temperature-dependent Ca^2+^ signals to transduce into the male sex determination pathway in turtles (Li et al., 2025; Ye et al., 2025). Here, using ricefield eel as a model we showed that a portion of Trpv4-expressing granulosa cells in ovary are responding to temperature cues, and initiates the sex reversal by increasing the expression of male pathway genes. This is supported by that Dmrt1 expression showed up within follicles in a typical granulosa cell location (Figure 3E; 6C). We propose that warm temperature exposure may cell-autonomously reprogram of a portion of Trpv4^+^ granulosa cells into Dmrt1/Sox9a-positive Sertoli precursor cells, thereby promoting the sex change of ricefield eel.

To summary, this study used ricefield eel hermaphrodite to elucidate the molecular basis underlying the transduction of environmental temperature into an intracellular signal for sex determination. We have made a few important findings. First, the ovarian somatic cells in ricefield eel are highly responding to the warm temperature. Warm temperature is sufficient to induce the expression of male pathway genes, without input of other factors such as hormones. Second, we identified a group of Trpv4-expressing granulosa cells that can directly perceive and respond to ambient temperature changes. Third, the pStat3-Kdm6b-*dmrt1* axis is downstream and mediates temperature-evoked Trpv4 activation. Our work revealed a comprehensive TISR mechanism, which involves signals that initiate sex reversal (temperature) and the capture (Trpv4), sensing (Trpv4-controlled calcium influx), transduction and interpretation (the Jak/Stat3 pathway) of environmental signals into the sex determination pathway (*kdm6b/dmrt1*). Of course, this work has limitations. Due to the unique life history of ricefield eel, genetic evidences are not sufficiently provided to substantiate the conclusion. Direct evidences for Trpv4 control of Ca^2+^ signaling are lacking.

## ACKNOWLEDGEMENT

This work was supported by the National Key Research and Development Program of China (2022YFD2400101) to YH Sun. We thank Tanhong Eel Industry Aquaculture Co., Ltd (Chibi city, Hubei Province) for collecting the fish materials and fish framing. We thank Yue Ou from Sun’lab for drawing the schematic role of Trpv4. We are grateful to Prof. Jianzhen Li from Northwest Normal University for critical comments for this work.

## AUTHOR CONTRIBUTIONS

Z Yang, TT Luo and YM Zhang performed experiments and generated data; YH Sun designed experiments, supervised the work, and wrote the manuscript; all authors edited the manuscript.

## RESOURCE AVAILABILITY

### Lead contact

Further information and requests for resources and reagents should be directed to and will be fulfilled by the lead contact, Yuhua Sun.

### Materials availability

All antibodies and plasmids generated in this study are available from the lead contact.

## Materials and Methods

### Sampling and maintenance of ricefield eel

Ricefield eel were purchased from the Baishazhou market, Wuhan, China. They were temporarily maintained in the lab at 25±1 °C under a 14–h light/10–h dark cycle, and fed daily with commercial diet. Female fish were usually less than 40 cm in length, males were more than 50 cm in length, and intersex animals were of medium length.

Animal experiments and treatments were performed according to the Guide for Animal Care and Use Committee of the Institute of Hydrobiology, Chinese Academy of Sciences (IHB, CAS, Protocol No. 2016-018).

### RNA preparation and RNA-sequencing

We have performed RNA-sequencing (RNA-seq) experiments using gonadal tissues from female, middle-late intersex, and male animals (Zhang et al., 2025). To explore the earliest events that trigger the onset of sex change of ricefield eel, gonads from female, early intersex, and middle intersex animals were isolated. Each gonad was examined by color and morphology, and in some cases histological experiments were performed to confirm the gonadal identities. Two to three gonads from each group were pooled, and total RNAs were isolated using a TRIzol reagent (ThermoScientific, USA). RNA sample quality was checked by the OD 260/280 ratio using a Nanodrop 2000. The total mRNAs were sent to the BGI (Beijing Genomics Institute) company (ShenZhen, China), where RNA-sequencing libraries were constructed and sequenced by a BGI-500 system. RNA-seq experiments were performed at least two times, with two technical repeats.

### Quantitative real-time PCR (qPCR) experiments

Total RNA was isolated using the Isolation Kit mRNA. *dmrt1/sox9a* were used as male-specific genes, and *cyp19a1a/foxl2* were used as female-specific genes. The primers and the related information used in this work were listed in Table 2. qPCR analysis was used to determine gene expression levels. A total of 1 µg RNA was reverse transcribed into cDNAs using the TransScript All-in-One First-Strand cDNA synthesis Supermix (Transgen Biotech, China, AT341). qPCR amplification was carried out on a Bio-Rad CFX96 Touch Real-Time PCR System (Bio-Rad, Hercules, CA, USA) in triplicate. The reaction mixture consisted of 5 µL PerfectStart™ Green qPCR SuperMix (Transgen Biotech, China), 3.6 µL ddH2O, 0.2 µL forward and reverse primers, and 1 µL cDNA. The cycling parameters used were 94 °C for 30 s, 94 °C for 5 s, 60 °C for 15 s, and 72 °C for 10 s for 40 cycles. Quantification cycle or cycle threshold values were determined using CFX Manager 3.1 (Bio-Rad, USA). Primers were 20-21 nucleotides long, with a melting temperature between 58 and 60 °C and a guanine-cytosine content between 50% and 60% generating an amplicon of 80-200 bp. Beta-actin was used as a reference gene. All qPCR experiments were repeated three times, and the relative gene expression levels were calculated based on the 2^−ΔΔCt^ method.

### siRNA knockdown experiments

To study in vivo function of Trpv4, we used siRNA to deplete the expression of the *trpv4* gene. The sequences of the ricefield eel *trpv4* gene (Accession Number: NW_018127903.1) were obtained from NCBI GenBank. The siRNA sequences are list in Table S2. These siRNAs were purchased from Sangon Biotech (Shanghai, China). Eighteen female fish were equally divided into 3 groups: 25 □+ MOCK; 34 □+ MOCK; 34 □+ *trpv4*-siRNA. For the 34 □-group setting, after 1-2 day of acclimation at 25□, the temperature of water was gradually increased by 3□ per day until reaching to 34□. For RNAi experiment, before increasing the temperatures, the siRNA was injected into ovaries through genital papilla. siRNAs were injected twice, for an interval of 2 days. Three *trpv4*-siRNAs were mixed in equal amounts; each fish was injected with 100 nM/kg. The MOCK groups were injected with equal amounts of control siRNAs. Two days after the second siRNA injection, ovarian samples were processed for qPCR analysis, and/or cryopreserved for ISH.

### WB analysis

WB was performed as previously described (Sun et al., 2020; 2023). The antibodies used in this work were: Amh (Huabio, #HA500137, China), Dmrt1 (home-made), Sox9a (home-made), Foxl2 (Thermofisher, #PA1-802, USA), Stat3 (Cell signaling, #9139, USA), pStat3 (Cell Signaling, #9145, USA). To validate the specificity of the antibodies, siRNA-mediated knockdown with immunoblot quantification with at least 2 replicates were performed (Supplementary File 1).

### IF experiments

Anesthetic fish were fixed with 4% PFA, and gonads were isolated. The gonads were washed three times with PBS and dehydrated in sucrose solution (15% sucrose/PBS, 30% sucrose/PBS) for 2 h at 4 □. Gonads were mounted in Tissue-Tek OCT compound (#4583, Sakura) and sectioned to 30 µm thickness on a cryostat. Slides containing sections were dried at room temperature for 30 min and washed 3 times for 5 min at room temperature with PBS. Sections on slides were blocked using 5% normal bovine serum (#A2153, Sigma Life Science) in PBS + 0.1% Triton-X100 (#V900502, VETEC) for 2 h. After wash, Primary antibodies, including Amh (Huabio, China), Foxl2 (ab5096, abcam), Trpv4 (Huabio, ER65407, China), Dmrt1 (home-made), Sox9a (home-made), Vimentin (OMA1-06001, Thermofisher, USA), were added at dilution of 1: 1000. Slides were washed 3 times with PBS for 5 min, followed by incubation in Alexa 488 or Alexa 555 (#A11008/A21428, Thermofisher, USA) secondary antibodies (1: 500) for 2 h. The samples were counter-stained with DAPI (#D9542, Sigma, 1:1000) in 1×PBS at room temperature for 1 h. After 3 times washing with PBS, slides were mounted using an anti-fade mounting medium (#HY-K1042, MedChemExpress, China). Mounted slides were imaged with a Leica Confocal Microscope (TCS SP8 STED, Germany).

### In situ hybridization (ISH)

To detect the expression of *trpv4/kdm6b* in the gonads, ISH experiments were performed. The cDNAs of ricefield eel *trpv4* and *kdm6b* were amplified by gene specific primers *trpv4*-SP6-F: ATTTAGGTGACACTATAGAAGCGTTTCTAGCCATTTCCTATCGT, and *trpv4*-T7-R: TAATACGACTCACTATAGGGAGACATTATCTGCTCCTAATCGAACC. All other primers can be found in Table S1. Digoxin labeled RNA probes of Sp6-sense and T7-antisense were synthesized using the DIG RNA Labeling Kit Sp6/T7 (Roche, Basel, Switzerland). ISH was conducted following the methodology outlined below. Briefly, the fixed gonads were processed by dehydration, paraffin embedding and serial sectioning (5 µm). Then the gonad slices were digested at 37 □ with 200 ng/mL proteinase K for 5 min. Hybridization was carried out for 16 h at 60 □ using a probe concentration of 1 ng/µL in the hybridization buffer. The samples were incubated with the Anti-Digoxigenin-AP conjugate (Roche, Basel, Switzerland) at a 1: 2500 dilution for 16h at 4 □, and stained in NBT/BCIP staining solution (Roche, Basel, Switzerland) in the dark for 0.5–1.5h at room temperature. The results were observed and photographed using an optical microscope (Zeiss, Oberkochen, Germany). Drawings and final panels were designed using Adobe Photoshop CS6 (San Jose, CA, USA).

### ChIP experiments

ChIP experiments were performed according to the Agilent Mammalian ChIP-on-chip manual. Briefly, gonadal tissues were processed into single cells and were fixed with 1% formaldehyde for 10 min at room temperature. The reactions were stopped by 0.125 M Glycine for 5 min with rotating. The fixed chromatin was sonicated to an average of (500-1000) bp (for ChIP qPCR) using the S2 Covaris Sonication System (USA) according to the manual. Then Triton X-100 was added to the sonicated chromatin solutions to a final concentration of 0.1%. After centrifugation, 50 µL of supernatants were saved as input. The remainder of the chromatin solution was incubated with Dynabeads previously coupled with 5 µg ChIP grade pStat3 antibodies overnight at 4 °C with rotation. The next day, after 7 times washing with the wash buffer, the complexes were reverse cross-linked overnight at 65 °C. DNAs were extracted by hydroxybenzene-chloroform-isoamyl alcohol and purified by a Phase Lock Gel (Tiangen, China). The ChIPed DNAs were dissolved in 100 µL distilled water. qPCR was performed using a Bio-Rad instrument. The enrichment was calculated relative to the amount of input as described. All experiments were repeated at least two times. The relative gene expression levels were calculated based on the 2^−ΔΔCt^ method. The paired t-test was used for the statistical analysis. Data were shown as means± SD.

### Luciferase Assay

HEK293T cells were seeded in 24-well plates in DMEM medium containing 10% FBS for 24h. The cells were then transiently transfected with the pGL4-*kdm6b*-luc or pGL4-*kdm6b*M-luc reporters using Lipofectamine 2000 (Invitrogen). pTKRenilla was used as an internal control. The luciferase activity was measured with the Dual-luciferase Reporter Assay system (Promega).

### Hematoxylin and eosin (H&E) experiments

H&E experiments were used for the identification of gonadal types in ricefield eels. The gonads were fixed in Bouin’s solution for at least 24 h, and the H&E experiments were performed by Wuhan Icongene Biotechnology Company. Briefly, dehydration and paraffin imbedding were then performed on the ASP6025S Automatic Vacuum Tissue Processor (Leica, Wetzlar, Germany). The samples were sectioned using the Leica microtome (Leica) at a thickness of 5 µm. After de-paraffinization, hydration and staining, the sections were examined on the Nikon ECLIPSE Ni-U microscope and micrographs were taken with the Digit Sight DS-Fi2 digital camera (Nikon).

### Gonadal sex identification

The gonadal types were initially identified according to morphological features including the size, shape and color. The gonadal sex of each fish was confirmed by histological sectioning and microscopic observation. In some cases, gene expression analysis was also used to confirm the gonadal sex types. Male genes such as *dmrt1* were not expressed in ovaries, slightly up-regulated in early ovotestes, and abundantly expressed in middle- and late-ovotestes.

### Long term temperature experiments

The aim of this experiment was to assess the gonadal phenotypes of female fish that were reared at 25 °C (cool temperature, CT) vs 33/34 °C (warm temperatures, WT) over 6 months. The experiments were performed from September–3, 2024. 1.5–year–old wild female fish were transiently maintained for 3-5 day at the laboratory, and unhealthy animals were discarded. A total of approximately 400 female fish were randomly divided into the CT and WT groups, stocked in 10–12 tanks, at a density of 15–20 fish per tank. Fish were fed with commercial diet.

1, 3, 6 months later, one tank in each group was randomly selected, and fish were anaesthetized and measured for body length and weight. The gonads were isolated and subjected to the histological analysis to determine the gonadal sex types. In some cases, gene expression analysis was used to determine the gonadal sex.

### Short term temperature experiments

#### Animal experiments

The experiments were used to explore how warm temperature treatment affects the expression of sex differentiation genes in ovaries in a short period of time (3–10 days). The ricefield eels were reared at 25 °C (cool temperature) and at 34 °C (warm temperature). The temperature–increase protocol started at 25 °C, with a progressive increase of 3 °C per day until reaching 34 °C. Temperature was monitored twice a day throughout the experiment. Fish were fed with Artemia daily.

The ricefield eels from the Baishazhou market were transiently raised for 1–2 day at cool temperature (25 °C). The fish were then divided into 4 groups based on the temperature and the injected small molecules: 25 °C+ DMSO; 25 °C+ GSK1016790A; 34 °C+ DMSO; 34 °C+ RN1734. For the 34 °C group setting, the temperature of water was gradually increased by 3 °C every day until reaching to 34 °C. Before increasing the temperatures, the small molecules of appropriate doses were injected into ovaries. The final concentrations used were at 0.02 mg/kg body weight for RN1734, 0.01 mg/kg body weight for HO-3867 or GSK1016790A or Colivelin, and a similar volume of 1‰ DMSO were injected and served as control.

To determine the upstream and downstream relationships between Trpv4 and pStat3, rescue experiments were performed by injecting the small molecules into the ovaries. Six groups were set up based on the temperature and the injected small molecules: 25 °C+ DMSO; 25 °C+ GSK1016790A; 25 °C+ GSK1016790A+ HO3867; 34 °C+ DMSO; 34 °C+ RN1734; 34 °C+ RN1734+ Colivelin.

To investigate the role of Kdm6b, 0.5 µM GSK-J4, an Kdm6b specific inhibitor, was injected into the ovaries or added into the cultured ovarian explants and/or cells.

#### Ovarian explant or cell culture

The ovaries were isolated from female ricefields and washed with cold 2% pen/strep PBS 3 times. The ovaries were cut into 2 mm^3^ pieces, and/or were digested by 0.25% TrypE for 30 min into single cells. For single cell culture, after filtration, ovarian cells were plated and cultured in 12 well plates.

For pharmaceutical experiments for Trpv4, the cells were divided into 4 groups: 26 °C+ DMSO; 26 °C+ GSK1016790A; 33 °C+ DMSO; 33 °C+ RN1734. For the 33 °C group setting, after 1–2 day of acclimation at 25 °C, the temperature of water was gradually increased by 3 °C every day until reaching to 33 °C. The doses of small molecules were optimized, and the final concentrations used were at 10 μM for RN1734, 100 nM for GSK1016790A and a similar volume of 1‰ DMSO were added and served as control. For pharmaceutical experiments for pStat3 function, 2 μM HO3867 and 20 μM Colivelin were added.

To determine the upstream and downstream relationships between Trpv4 and pStat3, rescue experiments were performed. Six groups were set up based on the temperature and the small molecules: 25 °C+ DMSO; 25 °C+ GSK1016790A; 25 °C+ GSK1016790A+ HO3867; 33 °C+ DMSO; 33 °C+ RN1734; 33 °C+ RN1734+ Colivelin.

#### Statistic analysis

For gene expression analyses, differences in mean values between two groups were assessed using Student’s t-test. A one-way ANOVA was used to compare the expression levels of each target and the differences were determined using Tukey’s post hoc test. Significance was defined as \**p* <0.05, \*\**p* <0.01, \*\*\**p* <0.001, \*\*\*\**p* <0.0001. Data are presented as mean standard error (SEM). Statistical analyses were conducted and graphs.

## Supplementary Figure legends

**Figure 1-figure supplement 1.**
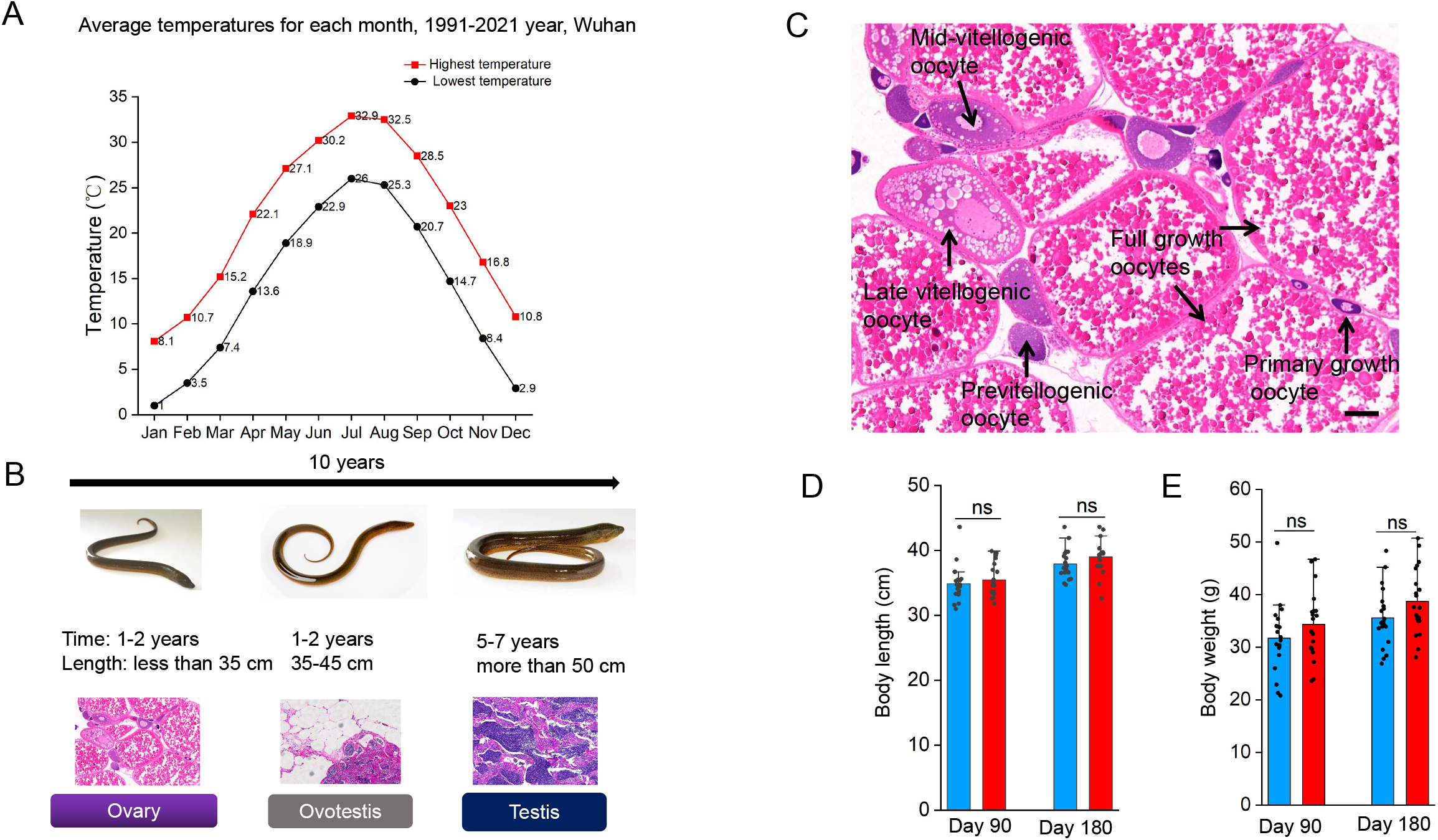
(A) The graph showing the annual temperature dynamic by month for 30 years in Wuhan area, Hubei province, China. The highest and lowest temperatures per month were shown. (B) Schematic of the whole process of sex change, summarizing morphology, length and gonadal histology across time (10 years). The ricefield eel are born as females. After 2 year growth, they reach sexual maturity. After spawning (ranging from May to August each year), females will enter 1-2 years of intersex stage before becoming functional males. The average lifespan of wild ricefield eel is around 10 years. (C) H&E staining images showing the typical cell types in an ovary of a 2-year-old female, before spawning. Bar: 200 µm. (D) The average body length of fish that were raised for 90 and 180 days at the indicated temperatures. (E) The average body weight of fish that were raised for 90 and 180 days at the indicated temperatures. ns: not significant. The experiments were repeated two times.

**Figure 2-figure supplement 1.**
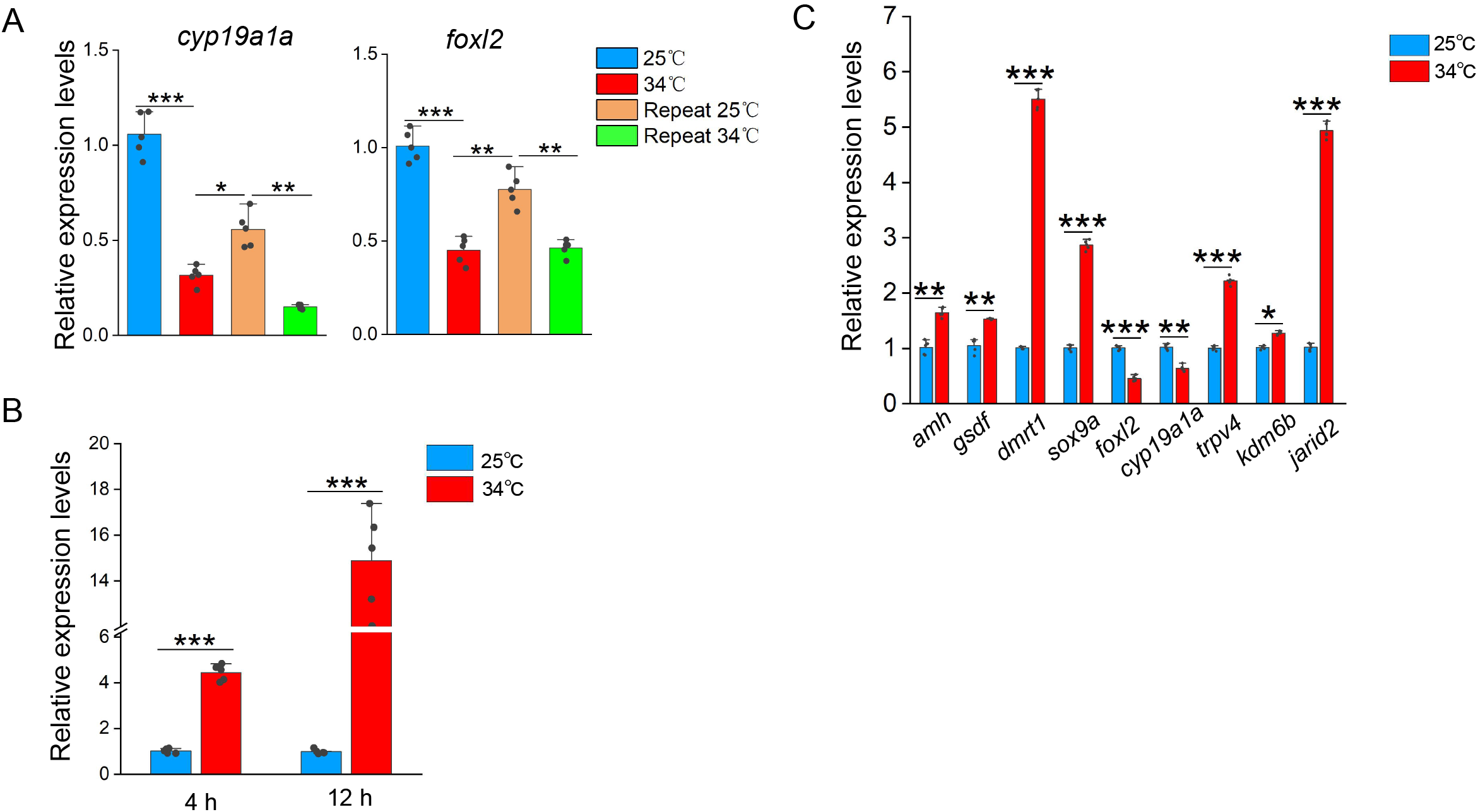
(A) The qPCR results showing the expression patterns of the indicated female sex genes in repeated temperature shifting experiments of in vitro cultured ovaries. n=5 per group. (B) The qPCR results showing the expression of *trpv4* at the indicated time points of in vitro cultured ovarian explants. n=5 per group. (C) The qPCR results showing the expression of the indicated temperature responding- and sex-biased genes in ovaries of females that reared at 25 □ and after 1 day exposure to 34 □. n=5 per group. *: *P*< 0.05, **: *P*< 0.01, ***: *P*< 0.001, and ****: *P*< 0.0001. ns: not significant. All experiments were repeated at least three times.

**Figure 3-figure supplement 1.**
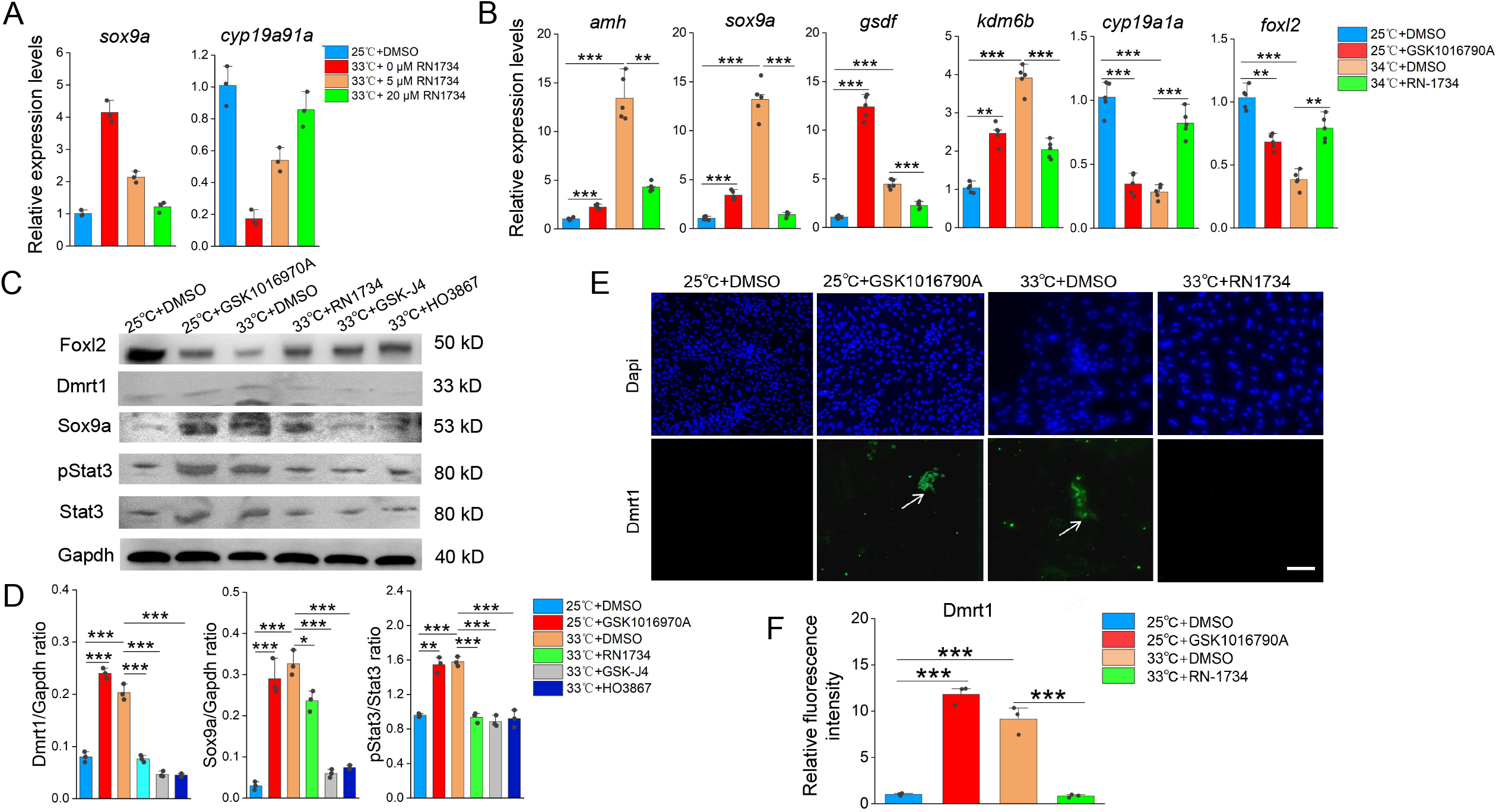
(A) qPCR results showing the expression of the indicated male- and female-biased genes in ovarian explants cultured at 25 □ and 34 □ with increasing doses of small molecule RN1734. n=5 per group. (B) qPCR results showing the expression of the sex-biased genes at the indicated conditions, based on in vitro cultured ovarian explants. n=5 per group. (C) WB images showing the expression levels of the indicated proteins at the indicated conditions, based on in vitro cultured ovarian explants. (D) Quantification of panel C. (E) IF images showing the expression levels of Dmrt1 at the indicated conditions, based on in vitro cultured ovarian explants. n=5 per group. (F) Quantification of panel C. *: *P*< 0.05, **: *P*< 0.01, ***: *P*< 0.001, and ****: *P*< 0.0001. All experiments were repeated at least three times.

**Figure 5-figure supplement 1.**
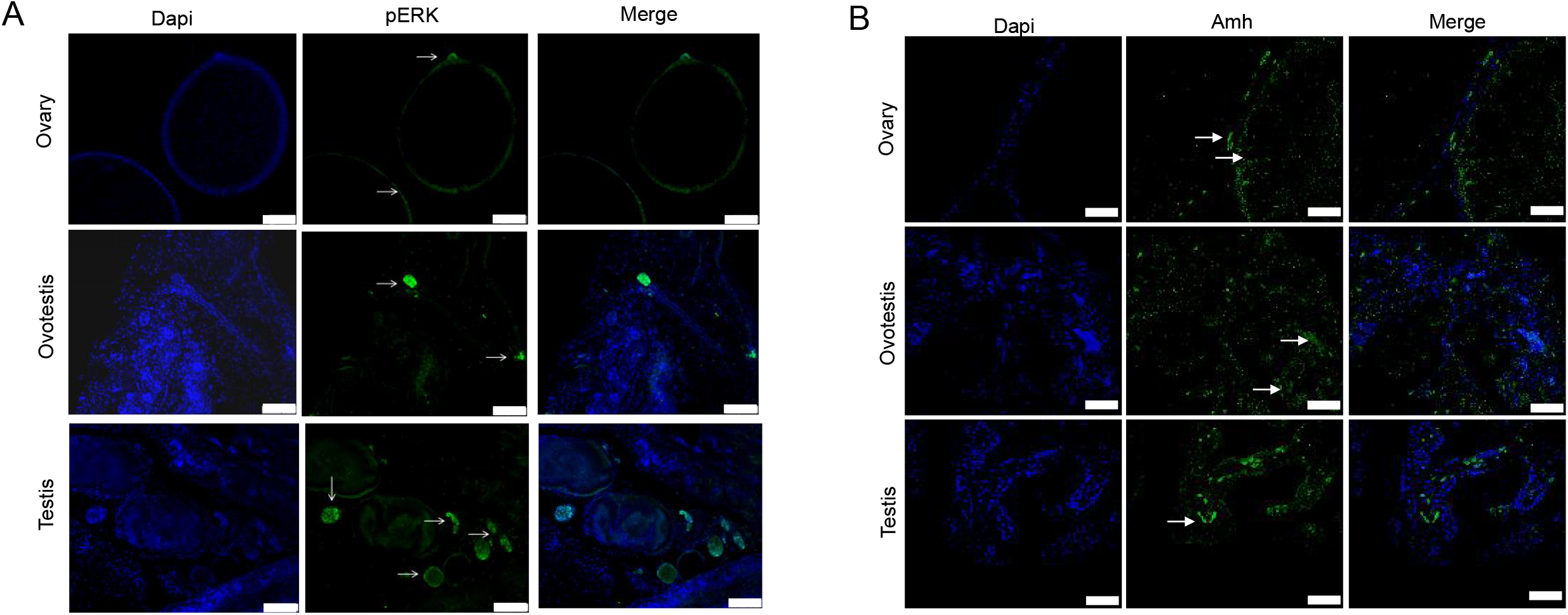
(A-B) The representative IF images showing the expression of pERK and Amh from ovaries, ovotestes, and testes. n=6 per group. The experiments were repeated at least two times.

**Figure 6-figure supplement 1.**
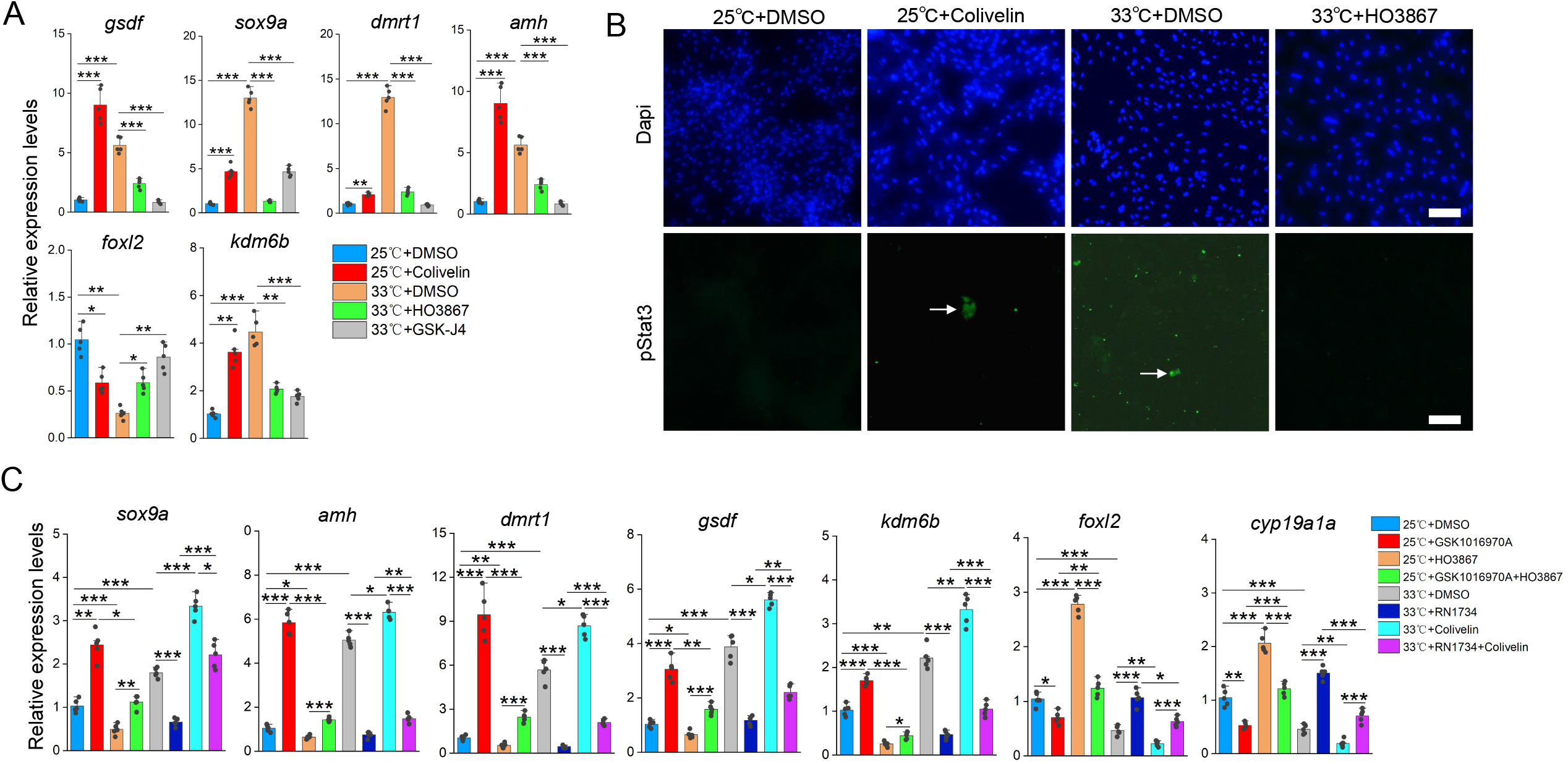
(A) qPCR results showing the expression of the indicated genes at the indicated conditions, based on in vitro cultured ovarian explants. (B) Representative IF images showing the expression of pStat3 at the indicated conditions, based on in vitro cultured ovarian explants. n=6 per group. (C) qPCR results showing the expression of the indicated genes at the indicated conditions, based on in vitro cultured ovarian explants. *: *P*< 0.05, **: *P*< 0.01, ***: *P*< 0.001, and ****: *P*< 0.0001. ns: not significant. All experiments were repeated at least three times.

## REFERENCES

Castelli, M.A., Whiteley, S.L., Georges, A., Holleley, C.E., 2020. Cellular calcium and redox regulation: the mediator of vertebrate environmental sex determination? Biol. Rev. Camb. Phil. Soc. 95, 680–695. 10.1111/brv.12582.

Chan, S.H, Phillips, J.G., 1967. Structure of gonad during natural sex reversal in Monopterus albus (Pisces: Teleostei). Journal of Zoology, 151(1): 129–141.

Chen, Q., Yang, D., Chen, M., Xiong, J., Huang, J., Ding, W., Gao, K., Lai, B., Zheng, L., Tang, Z., Zhang, M., Yan, T., He, Z., 2024. Smad4 and FoxH1 potentially interact to regulate cyp19a1a promoter in the ovary of ricefield eel (Monopterus albus). Biol. SexDiffer. 15, 60–71

Cheng, H., Zhou, R., 2022. Decoding genome recombination and sex reversal. Trends Endocrinol. Metabol. 33 (3), 175–185.

Deveson, I.W., Holleley, C.E., Blackburn, J., Marshall Graves, J.A., Mattick, J.S., Waters, P.D., Georges, A., 2017. Differential intron retention in Jumonji chromatin modifier genes is implicated in reptile temperature-dependent sex determination. Sci. Adv. 3, e1700731, 1700739.

Dupont, S., Garcia-Moreno, A., V O’Neil E., Capel B., 2025. KDM6B is a conserved activator at the top of the male sex determination pathway. Development. 152(12):dev204724.

Fan, M., Yang, W., Zhang, W., Zhang, L., 2022. The ontogenic gonadal transcriptomes provide insights into sex change in the ricefield eel Monopterus albus. BMC Zool. 7, 56–71.

Fan, M., Yang, W., Sun, S., Li, Z., Zhang, L.H., Zhang, W., 2021. Natural sex change of the”virgin”female ricefield eel Monopterus albus, 45: 387–396. doi: 10.7541/2021.2020.142.

Ge, C.T., Ye, J., Weber, C., Sun, W., Zhang, H.Y., Zhou, Y.J., Cai, C., Qian, G.Y., Capel, B., 2018. The histone demethylase KDM6B regulates temperature-dependent sex determination in a turtle species. Science, 360: 645–650.

Güler, A.D., Lee, H., Iida, T., Shimizu, I., Tominaga, M., Caterina, M., 2002. Heat-evoked activation of the ion channel TRPV4. J Neurosci. 22, 6408–6414.

Haltenhof, T., De Bortoli, F., Schiefer, S., Meinke, S., Emmeriches A.K., Petermann, K., Timmermann, B., Imhof, P., Franz, A., Loll, B., Wahl, M.C., Preushner, M., Heyd, F., 2020. A conserved kinase-based body-temperature sensor globally controls alternative splicing and gene expression. Molecular Cell, 78: 57–69.

He, Y., Shang, X., Sun, J., Zhang, L., Zhao, W., Tian, Y., Cheng, H., Zhou, R., 2010. Gonadal apoptosis during sex reversal of the rice field eel: implications for an evolutionarily conserved role of the molecular chaperone heat shock protein 10. J. Exp. Zool. B Mol. Dev. Evol. 314, 257–266. 10.1002/jez.b.21333.

He, Z., Deng, F., Yang, D., He, Z., Hu, J., Ma, Z., Zhang, Q., He, J., Ye, L., Chen, H., He, L., Luo, J., Xiong, S., Luo, W., Yang, S., Gu, X., Yan, T., 2022a. Crosstalk between sexrelated genes and apoptosis signaling reveals molecular insights into sex change in a protogynous hermaphroditic teleost fish, ricefield eel Monopterus albus. Aquaculture 552, 737918–737931. 10.1016/j.aquaculture.2022.737918.

He, Z., Ye, L., Yang, D., Ma, Z., Deng, F., He, Z., Hu, J., Chen, H., Zheng, L., Pu, Y., Jiao, Y., Chen, Q., Gao, K., Xiong, J., Lai, B., Gu, X., Huang, X., Yang, S., Zhang, M., Yan, T., 2022b. Identification, characterization and functional analysis of gonadal long noncoding RNAs in a protogynous hermaphroditic teleost fish, the ricefield eel (Monopterus albus). BMC Genom. 23, 450–467.

He, Z., Xiao, F., Yang, D., Deng, F., Ding, W., He, Z., Wang, S., Chen, Q., Wang, H., Chen, M., Gao, K., Xiong, J., Tang, Z., Zhang, M., Yan, T., 2024. Protein expression patterns and metal metabolites in a protogynous hermaphrodite fish, the ricefield eel (Monopterus albus). BMC Genom. 25, 500–515

Holleley, C.E., O’Meally, D., Sarre, S.D., Graves, J.M., Ezaz, T., Matsubara, K., Azad. B., Zhang, X.W., Georges, A., 2015. Sex reversal triggers the rapid transition from genetic to temperature dependent sex. 523: 79–83.

Hu, Q.M., Lian, Z.T., Xia, X.P., Tian, H.F., Li, Z., 2022. Integrated chromatin accessibility and DNA methylation analysis to reveal the critical epigenetic modification and regulatory mechanism in gonadal differentiation of Monopterus albus. Biology of Sex Differences, 13: 73–88.

Huang, J.W., Wang X.Y., Gao X.Y., Liu Q.H., Li, J. 2024. Transient receptor potential (TRP) channels in Sebastes schlegelii: Genome-wide identification and ThermoTRP expression analysis under high-temperature. Gene. 910: 148317

Huang, X., Guo, Y.Q., Shui, Y., Gao, S., Yu, H.S., Cheng, H.H., Zhou, R.J., 2005. Multiple alternative splicing and differential expression of dmrt1 during gonad transformation of the ricefield eel. Biology of reproduction, 73: 1017–1024.

Ji, F.Y., Yu, Q.X., Liu, J.D. 2001. Micro-isolation of individual chromosome for gene localization in ricefield eel. Chinese Science Bulletin, 22: 1894. 戢 福 云, 余 其 兴, 刘 江 东, 2001 。显 微 分 离 黄 鳝 单 条 染 色 体 用 于 基 因 定 位。科 学 通 报, 22: 1894. DOI: 10.3321/j.issn:0023-074X.2001.22.012

Jiang, Y.J., Luo, H.R., Hou, M.X., Chen, J., Tao, B.B., Zhu, Z.Y., Song, Y.L., Hu, W., 2022. Aromatase inhibitor induces sex reversal in the protogynous hermaphroditic rice field eel (Monopterus albus). Aquaculture 551, 737960–737972.

Kumar, A. Regulation of TRP channels by steroids: Implications in physiology and diseases. 2015. General and comparative endocrinology. 220: 23–32.

Li, J.M., Ye, Y.Z., Zheng, D., Pei, Z.D., Du, W.G., Yang, S.L., 2025. Molecular basis for thermal switch of temperature-dependent sex determination in a turtle species. Science Bulletin 70: 3319–3323

Liem, K.F. 1963. Sex reversal as a natural process in the synbranchiform fish Monopterus albus. Copeia, 2: 303–312.

Lin, J.Q., Zhou, Q., Yang, H.Q., Fang, L.M., Tang, K.Y., Sun, L., Wan, Q.H., Fang, S.G., 2018. Molecular mechanism of temperature-dependent sex determination and differentiation in Chinese alligator revealed by developmental transcriptome profiling. Sci. Bull. 63, 209–212. 10.1016/j.scib.2018.01.004

Lin, Q.H., Mei, J., Li, Z., Zhang, H.M., Zhou, L., Gui, J.F., Distinct and cooperative roles of amh and dmrt1 in self-renewal and differentiation of male germ cells in zebrafish. Genetics, 2017, 207: 1007–1022.

Liu, C.K., 1944. Rudimentary hermaphroditism in the symbranchoid eel Monopterus javanensis. Sinensia, (15): 1–8.

Liu, J. K., Gu, G.Y. 1950. Morphological changes in the gonad of Monopterus during sex reversal. Science, 2(3): 91.

Liu, L, Guo, M, Lv, X, Wang, Z, Yang, J, Li, Y, Yu, F, Wen, X, Feng, L and Zhou, T. 2021. Role of Transient Receptor Potential Vanilloid 4 in Vascular Function. Front. Mol. Biosci. 8:677661. doi: 10.3389/fmolb.2021.67766.

Liu, X.Y., Wang, L.C. 1987. The relationship between sex and age, body length, and weight. Freshwater fisheries, 6: 12–15.

Lu, J., Huang, S., Wei, S., Cheng, J., Li, W., Fei, Y., et al. 2025. Heat inducible nuclear translocation of Kdm6bb drives temperature dependent sex reversal in Nile tilapia. PLoS Genet 21(4): e1011664. 10.1371/journal.pgen.1011664.

Luo, T.T, Zeng, G.X., Xia, C.C., Hu, Z.X., Xue, L.Z., Sun, Y.H., 2026. Surrogate production of ricefield eel sperm by germ cell transplantation in adult zig-zag eel. Water Biology and Security (in press).

Luo, ZY et al. 2023. TRPV4 activation mediates Ca^2+^ influx and facilitates infectious spleen and kidney necrosis virus (ISKNV) replication. Journal of Virology, 97: 1–14.

Martínez-Pacheco, M., Díaz-Barba, K., Perez-Molina, R., Marmolejo-Valencia, A., CollazoSaldana, P., Escobar-Rodríguez, M., Sanchez-Perez, M., Meneses-Acosta, A., MartínezRizo, A., Sanchez-Pacheco, A.U., Furlan-Magaril, M., Merchant-Larios, H., Cortez, D., 2024. Gene expression dynamics during temperature-dependent sex determination in a sea turtle. Dev. Biol. 514, 99–108.

Mei, J., Gui, J.F., 2015. Genetic basis and biotechnological manipulation of sexual dimorphism and sex determination in fish. Science China Life Sciences 58, 124–136.

Mo, X.Y., Pang, P.Y., Wang, Y.L., Jiang, D.X., Zhang, M.Y., Li, Y., Wang, P.Y., Geng, Q.Z., Xie, C., Du, H.N., Zhong, B., Li, D.D., Yao, J. 2022. Tyrosine phosphorylation tunes chemical and thermal sensitivity of TRPV2 ion channel. eLife 11: e78301. DOI: 10.7554/eLife.78301.

Mundt, N., Spehr, M., Lishko, P.V. 2018. TRPV4 is the temperature-sensitive ion channel of human sperm. eLife, 7: e35853. DOI: 10.7554/eLife.35853

Shao, C.W. et al. 2014. Epigenetic modification and inheritance in sexual reversal of fish. Genome Research, 24: 604–615.

Shi, J., Sheng, D.L., Guo, J., Zhou, F.Y., Wu, S.F., Tang, H.Y., 2024. Identification of BiP as a temperature sensor mediating temperature-induced germline sex reversal in C. elegans. The EMBO Journal, 43: 4020–4048

Song, Y.L., Luo, H.R., Tao, B.B., Chen, J., Luo, D.J., Chen, K.X., Zhou, W.Z., Hu, W., 2022. Research Progress on the genetic basis of natural sex reversal and varieties breeding in Monopterus albus, Acta Hydrobiologica Sinica, 46(8): 1205–1214 10.7541/2022.2020.238

Steinberg, X., Lespay-Rebolledo, C., Brauchi, S., 2014. A structural view of ligand dependent activation in thermoTRP channels. Front. Physiol. 5, 171–185.

Tao, Y. X., Lin, H. R., Van der kraak, G., Peter R.E., 1993. Hormonal induction of precocious sex reversal in the ricefield eel, Monopterus albus. Aquaculture, 118: 131–140.

Tian, H.F., Hu, Q.M., Li, Z., 2021. A high-quality de novo genome assembly of one swamp eel (Monopterus albus) strain with PacBio and Hi-C sequencing data. G3-Genes Genomes Genetics, 11(1): jkaa032.

Todd, E.V., Ortega-Recalde, O., Liu, H., Lamm, M.S., Rutherford, K.M., Cross, H., Black, M.A., Kardailsky, O., Graves, M., Hore T.A., Godwin, J.R., Gemmell, N.J. 2019. Stress, novel sex genes, and epigenetic reprogramming orchestrate socially controlled sex change. Science Advances, 5: eaaw7006.

Vrenken, K.S., Jalink, K., van Leeuwen, F.N., Middelbeek, J., 2016. Beyond ionconduction: channel-dependent and -independent roles of TRP channels during development and tissue homeostasis. Biochim. Biophys. Acta, Mol. Cell Res. 1863, 1436–1446.

Wang, X., Lai, F.L., Xiong, J., Zhu, W., Yuan, B.F., Cheng, H.H., Zhou, R.J., 2020. DNA methylation modification is associated with gonadal differentiation in Monopterus Albus. Cell Biosci, 10:129. 10.1186/s13578-020-00490-4.

Wang, X., Lai, F.L., Shang, D.T., Cheng, Y.B., Lan, T., Cheng, H.H., Zhou, R.J., 2021. Cellular fate of intersex differentiation. Cell Death and Disease, 12: 388–402.

Wang, W.B., Zeng, B.P., Luo, Y.S., Han, Q., Liu, L.G. 2008. The relationship between sex and body length and weight of ricefield eel in DongTing lake area. Journal of Hunan Agricultural University (Natural Sciences), 34:469–473. 王 文 彬, 曾 博 平, 罗 玉 双, 韩 庆, 刘 良 国。2008。 洞 庭 湖区 黄 鳝 性 别 与 体 长 和 体 重 的 关 系, 湖 南 农 业 大 学 学 报, 34:469-473

Wang and Zeng. 2006. Report on Monopterus albus from Hainan Infected with Pallisentis (Neosentis) celatus and Eustrongylides Species. Sichuan Journal of Zoology, 125: 13–20

Weber, C., Zhou, Y.J., Lee, J.G., Looger, L.L., Qian, G.Y., Ge, C.T., Capel, B., 2020. Temperature-dependent sex determination is mediated by pSTAT3 repression of kdm6b. Science, 368: 3030–306.

Whiteley, S.L., Weisbecker, V., Georges, A., Gauthier, A.R., Whitehead D.L., Holleley C.E., 2018. Developmental asynchrony and antagonism of sex determination pathways in a lizard with temperature-induced sex reversal. Sci. Rep. 8 :14892.

Whiteley, S.L., Holleley, C.E., Wagner, S., Blackburn, J., Deveson, I.W., Marshall Graves, J.A., Georges, A., 2021. Two transcriptionally distinct pathways drive female development in a reptile with both genetic and temperature dependent sex determination. PLoS Genet. 17, e1009465, 1009498.

Whiteley, S.L., Wagner, S., Holleley, C.E., Deveson, I.W., Marshall Graves, J.A., Georges, A., 2022. Truncated jarid2 and kdm6b transcripts are associated with temperature-induced sex reversal during development in a dragon lizard. Sci. Adv. 8, eabk0275.

Wu, P.F., Wang, X.F., Ge, C.T., Jin, L., Ding, Z.H., Liu, F., Zhang, J., Gao, F., Du, W.G., 2024. pSTAT3 activation of Foxl2 initiates the female pathway underlying temperature-dependent sex determination. PNAS, 121: e2401752121. doi: 10.1073/pnas.2401752121.

Xiong, Y., Wang, S., Gui, J.F. & Mei, J., 2020. Artificially induced sex-reversal leads to transition from genetic to temperature-dependent sex determination in fish species. Science China Life Science, 2020, 63(1):157–159.

Yao, Z.L., Fang, Q.F., Li, J.Y., Zhou, M., Du, S.J., Chen, H.J., Wang, H., Jiang, S.J., Wang, X., Zhao, Y., Ji, X.S., 2023. Alternative splicing of histone demethylase kdm6b mediates temperature-induced sex reversal in the Nile tilapia. Current Biology, 33: 5057–5070.

Yatsu, R., Miyagawa, S., Kohno, S., Saito, S., Lowers, R.H., Ogino, Y., Fukuta, N., Katsu, Y., Ohta, Y., Tominaga, M., Guillette Jr., L.J., Iguchi, T., 2015. TRPV4 associates environmental temperature and sex determination in the American alligator. Sci. Rep. 5: 18581–18591.

Ye, Y.Z., Li, J.M., Li, W.M., Fu, X.N., Zhang, J.J., Yang, S.L., Du, W.G., 2025. Transcriptional control of male-specific pathway in temperature-dependent sex determination. Science Bulletin, 10.1016/j.scib.2025.07.025

Yamamoto et al., 2024. Trpv4-mediated apoptosis of Leydig cells induced by high temperature regulates sperm development and motility in zebrafish. COMMUNICATIONS BIOLOGY, 7: 96, 10.1038/s42003-023-05740-y.

Zhang, Y.M., Luo, T.T., Sun, Y.H., 2025. A temperature-induced sex reversal mechanism in ricefield eels. Water Biology and Security, 10.1016/j.watbs.2025.100360.

Zhou, L., Gui, J.F., 2016. Jian-Kang Liu: a pioneer of sex determination studies in vertebrates. Protein Cell 7, 1–3.

Honeycutt et al. 2019. Warmer waters masculinize wild populations of a fish with temperature-dependent sex determination. Scientific Reports, 9: 6527. 10.1038/s41598-019-42944-x.

Gutzke, W.H.N. & Crews, D., 1988. Embryonic temperature determines adult sexuality in a reptile. Nature 332, 832–834.

